# Proposal and extensive test of a calibration protocol for crop phenology models

**DOI:** 10.1101/2022.06.08.495355

**Authors:** Daniel Wallach, Taru Palosuo, Peter Thorburn, Henrike Mielenz, Samuel Buis, Zvi Hochman, Emmanuelle Gourdain, Fety Andrianasolo, Benjamin Dumont, Roberto Ferrise, Thomas Gaiser, Cecile Garcia, Sebastian Gayler, Matthew Harrison, Santosh Hiremath, Heidi Horan, Gerrit Hoogenboom, Per-Erik Jansson, Qi Jing, Eric Justes, Kurt-Christian Kersebaum, Marie Launay, Elisabet Lewan, Ke Liu, Fasil Mequanint, Marco Moriondo, Claas Nendel, Gloria Padovan, Budong Qian, Niels Schütze, Diana-Maria Seserman, Vakhtang Shelia, Amir Souissi, Xenia Specka, Amit Kumar Srivastava, Giacomo Trombi, Tobias K.D. Weber, Lutz Weihermüller, Thomas Wöhling, Sabine J. Seidel

## Abstract

A major effect of environment on crops is through crop phenology, and therefore, the capacity to predict phenology for new environments is important. Mechanistic crop models are a major tool for such predictions, but calibration of crop phenology models is difficult and there is no consensus on the best approach. Here we propose an original, detailed approach, a protocol, for calibration of such models. The protocol covers all the steps in the calibration work-flow, namely choice of default parameter values, choice of objective function, choice of parameters to estimate from the data, calculation of optimal parameter values and diagnostics. The major innovation is in the choice of which parameters to estimate from the data, which combines expert knowledge and data-based model selection. First, almost additive parameters are identified and estimated. This should make bias (average difference between observed and simulated values) nearly zero. These are “obligatory” parameters, that will definitely be estimated. Then candidate parameters are identified, which are parameters likely to explain the remaining discrepancies between simulated and observed values. A candidate is only added to the list of parameters to estimate if it leads to a reduction in BIC (Bayesian Information Criterion), which is a model selection criterion. A second original aspect of the protocol is the specification of documentation for each stage of the protocol. The protocol was applied by 19 modeling teams to three data sets for wheat phenology. All teams first calibrated their model using their “usual” calibration approach, so it was possible to compare usual and protocol calibration. Evaluation of prediction error was based on data from sites and years not represented in the training data. Compared to usual calibration, calibration following the new protocol reduced the variability between modeling teams by 22% and significantly reduced prediction error.

## 1. Introduction

Plant phenology is a major aspect of plant response to environment, and a major determinant of plant response to climate change. This includes phenology of natural vegetation, which is a dominant aspect of plant ecology (Cleland et al. 2007) and has been shown to be affected by warming (Piao et al. 2019; Menzel et al. 2020; Stuble et al. 2021) and phenology of cultivated crops. For the latter, phenology must be taken into account for crop management (Sisheber et al. 2022), choice of cultivar or cultivar characteristics adapted to a particular region (Zhang et al. 2022) and for evaluating the impact of climate change on crop production (Rezaei et al. 2018). It is thus important to be able to predict phenology as a function of environment, in particular as a function of climate.

A number of mechanistic crop models have been developed, which include simulation of phenology, and such models are regularly used to evaluate management options (McNunn et al. 2019) or the effect of climate change on crops, including wheat (Asseng et al. 2013), rice, (Li et al. 2015), maize (Bassu et al. 2014) and others. Such models are particularly important for taking into account an increasing diversity of combinations of weather events (Webber et al. 2020).

Mechanistic models in general, and models used to simulate crop phenology in particular (we will refer to such models as crop phenology models, though they are usually embedded within more general crop models), are based on our understanding of the processes and their inter-linkages that drive the evolution of the system. This conceptual understanding usually builds on detailed experiments that study specific aspects of the system (e.g. Brisson et al., 2003 for the crop model STICS). The set of model equations is referred to as “model structure” (Tao et al. 2018).

In addition to model structure, simulation requires values for all the model parameters. In essentially all uses of crop models, the model is first calibrated using observed data that is related to the target population for which predictions are required, for example observations for the specific variety of interest and/or for the particular set of growing environments of interest. Calibration normally only concerns a fairly small subset of the parameters in a crop model, but is essentially always necessary because mechanistic models are only approximations, without universally valid parameter values (Fath and Jorgensen 2011; Wallach 2011).

There are therefore two main tracks to improvement of crop phenology models. The first is through improvement of model structure through improved understanding of the underlying processes, and the second is through improvement of the model parameters. For a fixed data set, improvement of model parameters implies improvement of model calibration, and that is the topic here.

Calibration of crop models is usually patterned on methods developed for regression models in statistics. However, the application of statistical methods to system models is not straightforward. Among the difficulties are the fact that system models often have multiple output variables that can be compared to observed results (e.g., dates of heading and dates of flowering for crop phenology models), that errors for different variables in the same environment are often correlated and that there are usually many parameters, often more than the number of data points available. While the details differ, these problems apply to essentially all system models, not only crop phenology models but also full crop models, hydrological models, ecology models and models in other fields. No doubt as a result, there are no widely accepted standard methods for calibration of system models. It has been found, for example, that there is a wide diversity of calibration approaches between modeling teams furnished with identical data, even between modeling teams using the same model structure (Confalonieri et al. 2016; Wallach et al. 2021a, b), where a modeling team is a group of people working together on or with a crop model.

Because of the importance of calibration and the lack of standard approaches for calibration, there have been many studies published that make recommendations as to how to calibrate crop models or system models in other fields. One type of study is model-specific, and identifies the most important parameters to estimate for a particular model (Ahuja and Ma, 2011). Other studies have focused on the methodology of identifying the most important parameters through sensitivity analysis (Khorashadi Zadeh et al. 2022), on the choice between frequentist and Bayesian paradigms (Gao et al. 2021), on the form of the objective function, or on the numerical algorithm for searching for the best parameter values (Rafiei et al. 2022). A recent study has shown that different modeling teams make different choices for all the steps of the calibration procedure, (Wallach et al., 2021c). That study showed that the modeling community is far from having a consensus on how to calibrate phenology models, and emphasized the need to treat calibration holistically, taking into account all aspects.

The purpose of this study was to define and test an original, detailed, comprehensive procedure for crop phenology model calibration that could be applied to a wide range of crop phenology models. We refer to this new procedure as a “protocol”, to emphasize that it contains detailed instructions for calibration. It builds on the recommendations in Wallach et al. (2021c) but goes beyond those more general recommendations, most importantly in proposing an original approach for choosing the parameters to estimate, which is arguably the most important calibration decision. A second major innovation of the protocol is the definition of documentation tables for each step of the calibration procedure, which can be used both for communication within a modeling team and to inform users of the calibrated model. We tested the protocol in a large multi-model ensemble study where the objective was to simulate wheat phenology in multiple environments (combinations of site and sowing date) in France and in Australia. Nineteen modeling teams participated, and were all able to implement the protocol, showing that the protocol, though detailed, is nonetheless sufficiently flexible to be applicable to a wide range of models. The protocol was tested in comparison with the “usual” calibration procedure of each modeling team, which was possible because each modeling team had previously calibrated their model using the same data as here. To our knowledge this is the first example of such a stringent test for a new calibration procedure. It provides a realistic test of whether the proposed protocol really improves calibration. The current study showed that the protocol reduced the variability between modeling teams compared to usual calibration approaches and, most importantly, that it significantly reduced prediction error compared to usual calibration approaches.

## 2. Materials and Methods

### Data sets

Three data sets for wheat phenology were used here, each with data from multiple environments, i.e. combinations of site, year and sowing date. The first two data sets were from cultivar trials in France, for the two winter wheat varieties Apache and Bermude (Wallach et al 2021a). For both cultivars there were the same 14 calibration environments (6 different sites, sowing years 2010, 2011, 2014-2016 but not every year was represented for every site) and 8 evaluation environments (5 different sites, sowing years 2012 and 2013). The calibration and evaluation subsets had neither site nor year in common, so the evaluation is a rigorous test of how well a modeling team can simulate phenology for out-of-sample environments. The observed data were days from sowing to beginning of stem elongation (BBCH30 on the BBCH scale, Meier 1997) and to middle of heading (BBCH 55).

The third data set had data for the spring wheat variety Janz from a multi-location multi-year multiple sowing date trial in Australia (Lawes et al. 2016; Wallach et al. 2021b). The calibration data were from four sites in 2010 and 2011, with three sowing dates per site (overall 24 environments). The evaluation data were from six sites in 2012, with three sowing dates per site (overall 18 environmets). Once again, the calibration and evaluation subsets had neither site nor year in common. In the original trials, BBCH development stage was observed once weekly in each environment. Based on those data, a graph of BBCH stage versus day was produced, and interpolation was used to obtain the day for each integer BBCH stage from the earliest to the latest recorded in each environment. Those dates were provided to the modeling teams.

### Modeling teams

Nineteen modeling teams, using 16 different model structures (Table 1), participated in this study, which was carried out within the Agricultural Model Intercomparison and Improvement Project (AgMIP; www.agmip.org). The modeling teams are identified only by a code (“M1”, “M2” etc.) without indicating which model structure they used, since it would be misleading to give the impression that the results are determined solely by model structure.

**Table 1:** List of model structures used by participating modeling teams.

| Model structure | Version(s) | References |
| --- | --- | --- |
| AgroC | May 2018 | (Herbst et al. 2008; Klosterhalfen et al. 2017) |
| APSIM | 7.8, 7.9, 7.10 | (Keating et al. 2003; Holzworth et al. 2014) |
| AquaCrop | 4.0 | (Vanuytrecht et al. 2014) |
| CERES-Wheat | DSSAT V4.7. | (Hoogenboom et al., 2019a, 2019b; Jones et al., 2003) |
| CoupModel | Version 5.4.4 | (P.-E. Jansson 2012; Senapati et al. 2016; Coucheney et al. 2018) |
| CROPSIM-Wheat | DSSAT V4.7 | (Hoogenboom et al., 2019a, 2019b; Jones et al., 2003) |
| Cropsyst | 3.04.08 | (Stockle et al. 2001) |
| HERMES | 4.27 | (Kersebaum, 2007; Kersebaum, 2011) |
| LINTUL | LINTUL5 | (Wolf 2012) |
| MONICA | 2.02 | (Nendel et al. 2011; Specka et al. 2015, 2019) |
| PANORAMIX | R version | (Gate 1995; Chatelin et al. 2005) |
| SPASS | Expert-N 5.0 | (Wang 1997) |
| CERES | Expert-N 5.0 | (Godwin et al.) |
| SSM-Wheat |  | (Soltani et al. 2013) |
| STICS | 8_5_0 | (Brisson et al. 2009; Coucheney et al. 2015) |
| WOFOST | 7.1.7 | (Boogaard et al. 1998) |

### Goodness-of-fit and evaluation of predictions

Goodness-of-fit refers to how well a calibrated model fits the data used for calibration. Prediction accuracy refers to how well a calibrated model simulates for environments different than those in the calibration data set,. Since the evaluation environments here are for sites and years not represented in the calibration data, the test of simulated values against the evaluation data truly reflects how well a model can predict for new environments.

For both goodness-of-fit and out-of-sample prediction, our basic evaluation metric is the sum of squared errors (SSE) and the related quantities mean squared error (MSE) and root mean squared error (RMSE), where

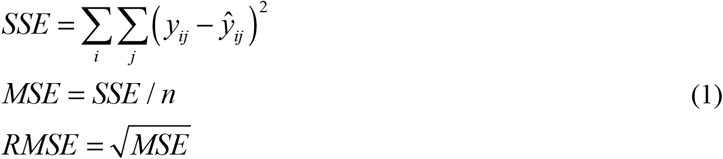

The sum for SSE is over variables and environments i.e. over all measurements for which there are corresponding simulated values. Here *y_y_* is the observed value of variable *i* for environment *j, ŷ_y_* is the corresponding simulated value and *n* is the number of terms in the sum. We also look at the decomposition of MSE as the sum of three terms, namely squared bias (bias^2^), a term related to the difference in standard deviations of the observed and simulated values (SDSD) and a term related to the correlation of observed and simulated vales (LCS) (Kobayashi 2004).

In addition, we compare the simulated results in this study with two simple benchmark models. The first (the “naive” model) is simply the average number of days to each stage in the calibration data of each data set. This is used as the prediction model for all environments of that data set. The often used Nash Sutcliffe modeling efficiency is one minus the ratio of MSE of a model to MSE of the naive model. The naive model ignores all variability between environments, so it is a very low bar as a benchmark. We therefore also use a more sophisticated benchmark, the “onlyT” model, introduced in Wallach et al. (2021a). This benchmark model assumes that the sum of degree days above a threshold of 0°C from sowing to each stage is fixed for spring wheat. For winter wheat, a simple vernalization model is used to determine the start of development. Vernalization is 0 if daily mean temperature is below −4°C, increases linearly to 1 at 3°C, remains at 1 to 10°C, decreases linearly to 0 at 17°C and is 0 above 17. °C. When the sum of daily vernalization reaches 50, vernalization is complete (van Bussel et al. 2015; Wallach et al. 2021a). Then the fixed number of degree days applies after vernalization is completed. Both benchmark models are quite easily parameterized based on calibration data, and then easily applied to new environments.

### Simulation exercise

The participants received input data (daily weather at the field, soil characteristics, management details and, where possible, initial conditions) for all environments of every data set. Also, the observed data from the calibration environments were provided to all participants. The participants were asked to use those data to calibrate their models using the calibration protocol described in detail below, and then to simulate and report days after sowing to stages BBCH10, BBCH30, and BBCH55 for the French calibration and evaluation environments, and to stages BBCH10, BBCH30, BBCH65, and BBCH90 for the Australian calibration and evaluation environments. Days to emergence (BBCH10) was included to have an example of a variable for which there were no calibration data. The BBCH stages 30 and 55 requested for the French environments represent stages that are used for fertilizer decisions in France. The BBCH stages 30, 65, and 90 requested for the Australian environments represent major transitions that are explicitly simulated by many models.

Seventeen of the 19 participating modeling teams participated in previous phases of this project, and in that framework had already calibrated their model with the same data as here using their usual calibration approach (Wallach et al. 2021a, b). The two remaining teams also calibrated their models using their usual approach before beginning to use the protocol proposed here. It is the results of the usual calibration method that are compared here to the results of using the proposed protocol. At no time were the evaluation data shown to participants, neither in previous studies nor in the present study.

The protocol does not impose a specific software solution. However, several participants used trial and error in their usual approach and requested help in finding and implementing an automated search algorithm, since that is required for the protocol. To answer this need, the CroptimizR R package (Buis et al. 2021) was modified to do the protocol calculations, and many of the participants used this software.

In addition to the individual models, we report on two ensemble models, created by taking the mean (the e-mean model) or the median (the e-median model) of the simulated values. These ensemble models were calculated both for the usual and protocol calibration results.

### AICc and BIC

The protocol prescribes a model selection criterion to decide which parameters to estimate. The corrected Akaike Information Crition (AICc) and the Bayesian Information Criterion (BIC) are two different criteria that are often used for model selection (Chakrabarti and Ghosh 2011). Both are based on model error, with a penalization term that increases with the number of estimated parameters. Assuming that model errors are normally distributed, the criteria are

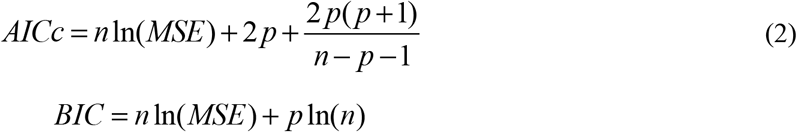

where *n* is the number of data points and *p* is the number of calibrated parameters. These criteria are only used for comparing models calibrated using the same data.

There have been comparisons between these criteria, but there does not seem to be one that systematically performs better than the other, for choosing the model that predicts best (Kuha 2004). In applying the protocol here, participants were asked to perform the calculations twice, once using the AICc criterion and once using the BIC criterion to choose the parameters to estimate. In almost all cases, the two criteria led to exactly the same choice of parameters. In the few cases where the criteria led to different choices, the final models had very similar RMSE for the evaluation data, with a very slight advantage to BIC (Supplementary tables S24-S25). All results shown here are based on the BIC criterion.

## 3. Results and discussion

### Description of protocol

The protocol is based on the recommendations in Wallach et al. (2021c), and follows the same list of steps (Figure 1), but has important additions, in particular for the choice of parameters to estimate and the documentation to be produced,

**Figure 1.**
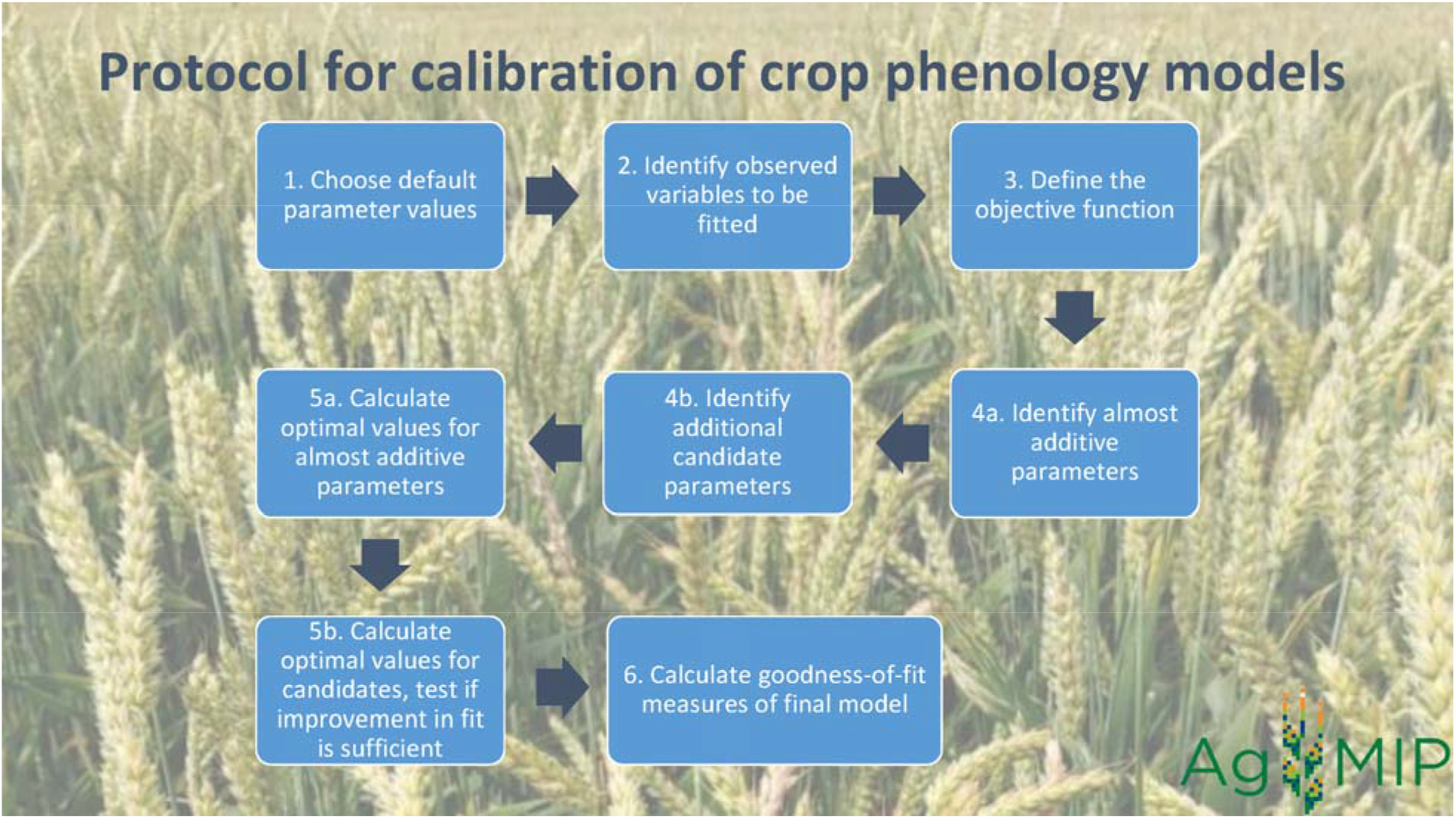
Schematic diagram of steps in proposed calibration protocol.

### Step 1. Choose default parameter values

Since most parameters retain their default values, the choice of default values for those parameters that affect phenology is important. For phenology, one would want to have reasonable approximations to the cycle length for the cultivar in question, to photoperiod dependence and to vernalization requirements. This information and more is usually available from the cultivar developer. The documentation for step 1 (see example in Table 2) specifies the characteristics of the cultivar being modeled and those of the cultivar used to provide default parameter values.

**Table 2:** Example of protocol documentation for Step 1, “Choose default parameter values”. The first row shows cultivar characteristics of the observed cultivar. The second row shows characteristics of the cultivar that provides the default parameter values. This example is for the French data set and modeling team M21.

| Variety | Characteristics |
| --- | --- |
| Variety of target population: Apache | A soft winter wheat. Stem elongation – semi-early. Heading – early. Vernalization requires 40 days where full vernalization occurs if daily average temperature is between 3°C and 10°C. There is no vernalization below -4°C or above 17°C. Otherwise there is a proportional reduction in vernalization effectiveness. |
| Default variety : Soissons | Soissons seems to be close to Apache in terms of vernalization requirements and earliness. |

### Step 2 Identify correspondence between observed and simulated variables

In the simplest case, there is a simulated variable that corresponds directly to each observed development stage. The documentation for step 2 is a table with one row for each observed variable, showing the corresponding simulated variable if any (see example in Table 3).

**Table 3:** Example of documentation for protocol Step 2, “Identify correspondence between observed and simulated variables”. The table has one row for each measured variable, showing the corresponding simulated variable if any. This example is for the French data sets and modeling team M21.

| Measured variable | Corresponding simulated value |
| --- | --- |
| Days to BBCH stage 30 | Days to end juvenile stage |
| Days to BBCH stage 55 | Days at maximum LAI |

### Step 3. Define the objective function

The objective function of the protocol is squared error summed over development stages and environments, which is the objective function of ordinary least squares (OLS) regression and is often used in crop model calibration. A major choice of the protocol is to include in the objective function all the observed development stages that have a simulated equivalent, including stages that are not of primary interest. A first reason is that often the same calibrated model will be used for several different objectives, so measured variables that are not of central interest in the current study may be important in future studies. Furthermore, using more variables makes the model a better representation of multiple aspects of the system dynamics, which is likely to improve all simulations. The choice in the protocol is to use OLS and to avoid estimating additional parameters related to variance and covariance of errors. However, one should check residual errors to evaluate the extent of heteroscedasticity or correlation of errors. Since the objective function is the sum of squared errors over the variables from step two, no new decisions are required here and no additional documentation is required.

### Step 4. Choose which parameters to estimate

This is arguably the most difficult, and the most important, decision of the calibration approach. Here we propose a novel approach which combines expert knowledge with a statistical model selection criterion. This approach distinguishes two categories of parameters to estimate: the nearly additive, obligatory parameters (those that will definitely be estimated) and the candidate parameters (those that will be tested, and only changed from the default value if the improvement in the fit to the calibration data is sufficiently large).

### Step 4a Identify the obligatory parameters

The obligatory parameters are parameters that are nearly additive, i.e. such that changing the parameter has a similar effect for all environments for some variable in the objective function. Usually, a parameter that represents degree days to a measured stage is a good choice as an obligatory parameter for time to that stage. Estimating a truly additive parameter (that adds the same constant amount to days to a stage for all environments) will exactly eliminate bias for that stage (i.e. the mean of simulated values will exactly equal the mean of observed values). Estimating an almost additive parameter will almost eliminate bias. Once bias is nearly eliminated, one may already have a fairly reasonable fit to the data. Each almost additive parameter must affect a different variable or combination of variables. There cannot be more almost additive parameters than the number of variables in the objective function. Otherwise, the parameters would be very poorly estimated, or non-estimable. The protocol does allow fewer almost additive parameters than observed variables. In that case bias is only nearly eliminated on average over several variables, and not for each variable. For each obligatory parameter one must provide the default value and what one considers a reasonable range for that parameter (for an example of choice of bounds see Tao et al. 2013). The documentation for step 4a is a table with one row for each obligatory parameter (see example in Table 4).

**Table 4:** Example of documentation for protocol step Step 4a, “Identify the obligatory parameters”. These are parameters that are almost additive, i.e. that have nearly the same effect for all environments. There is one row for each obligatory parameter. The number of obligatory parameters cannot exceed the number of observed variables which have a simulated equivalent, and each obligatory parameter must be nearly additive for a different variable or combination of variables. This example is for the French data set, variety Apache for modeling team M21.

| Name of obligatory parameter | explanation | Default value (lower-upper limits) |
| --- | --- | --- |
| stlevamf | Degree days sowing to end juvenile stage | 233 (150-400) |
| stamflax | Degree days sowing to maximum LAI | 354 (150-500) |

### Step 4b. Identify candidate parameters

The role of the candidate parameters is to reduce the variability between environments that remains after estimation of the obligatory parameters. It is the role of the modeler to identify the candidate parameters and to order them by amount of variability likely to be explained. In the calculation step (step 5), each candidate parameter is tested, and only those that lead to a reduction in the BIC criterion are retained for estimation. Otherwise, the parameter is kept at its default value. The documentation for step 4b is a table with one row for each candidate parameter (see example in Table 5).

**Table 5:** Example of documentation for protocol Step 4b, “Identify candidate parameters”. These are parameters that seem likely to explain a substantial part of the variability between environments that remains after fitting the obligatory parameters. There is one row for each candidate, which should be in the order of presumed importance. This example is for the French data set, variety Apache for modeling team M21.

| Candidate parameter (include units and a brief explanation) | Default value (Lower - upper bounds) |
| --- | --- |
| jvc (days): number of vernalizing days | 38 (25 – 60) |
| sensrsec (no unit): index of root sensitivity to drought (1=insensitive) | 0.5 (0 – 1) |
| belong (degree.day-1): parameter of the curve of coleoptile elongation | 0.012 (0.005 – 0.03) |
| JVCmini (days): minimum vernalizing days required | 7.0 (2 – 15) |
| stressdev (no unit): maximum phasic delay allowed due to stresses | 0 (0 – 1) |

### Step 5. Calculation of the optimal parameter values

The protocol prescribes the use of a simplex algorithm for searching for the optimal parameter values. The Nelder-Mead simplex method (Nelder and Mead 1965) is a robust, derivative-free method, which is appropriate for crop models which may have multiple discontinuities. The results of the simplex are sensitive to starting values (Wang and Shoup 2011), so the protocol calls for multiple starting points. In the first calculation step, the obligatory parameters are estimated and the BIC value is calculated. This is the initial list of parameter to estimate. Then each candidate parameter is tested in turn. If estimating the new candidate together with the previous list of parameters to estimate leads to a reduction in BIC, the candidate is added to the list of parameters to estimate. If not, the candidate returns to its default value and will not be estimated. The documentation for step 5 is a table with one row for each step in the calculation (see example in Table 6).

**Table 6:**
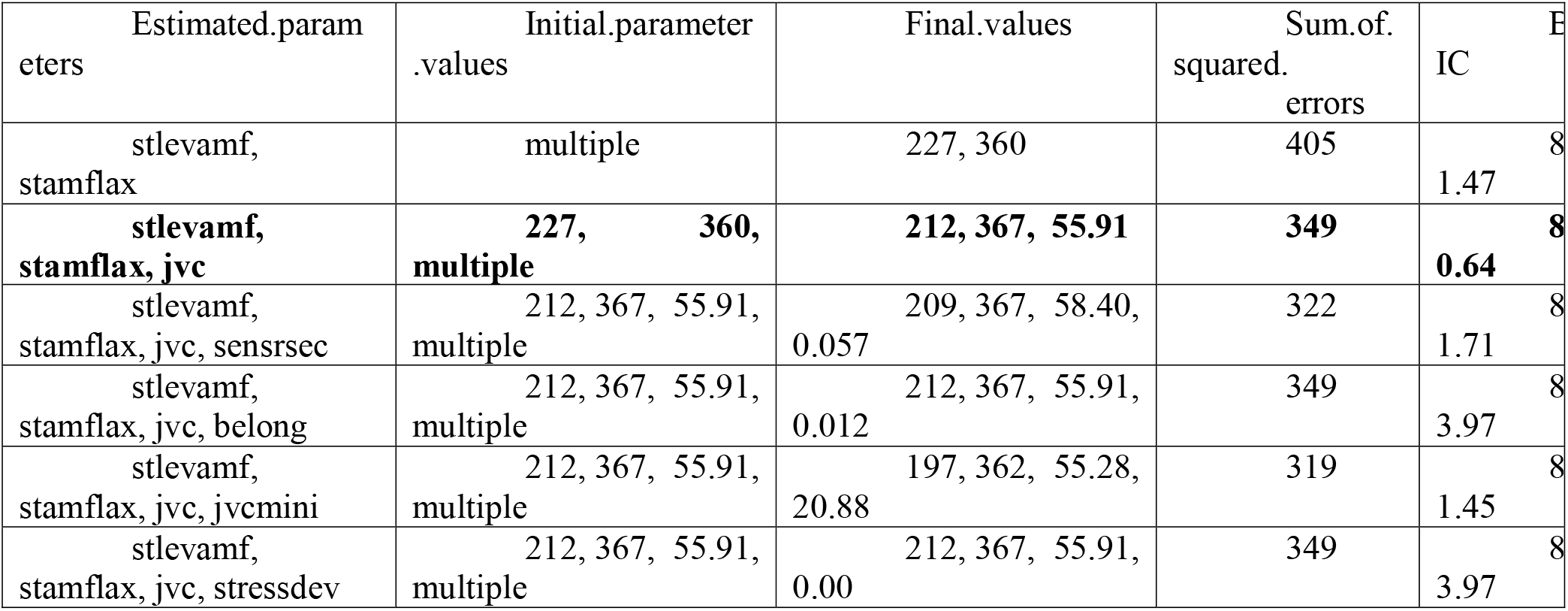
Example of documentation for Step 5 “Calculation of the optimal parameter values”. The first line shows the optimization results for the obligatory parameters, and the resulting sum of squared errors and BIC criterion. Each subsequent line corresponds to a candidate parameter. If estimating the candidate together with the previously selected parameters leads to a decrease in BIC compared to the smallest value so far, the candidate is added to the list of parameters to estimate. If not, the candidate returns to its default value and is not considered further. In this example, the first candidate parameter (jvc) is accepted, and all the subsequent candidate parameters increase BIC, and are therefore, rejected. The model finally chosen (minimum BIC) has three estimated parameters. This example is for the French data set, variety Apache, modeling team M21.

### Step 6 Examine goodness-of-fit

Many diagnostics are possible and useful. We emphasize particularly a graph of simulated versus observed values (see example in fig 2), calculated MSE and its decomposition for each variable (see example in table 7) and comparison with the two benchmark models (see example in table 8).

**Figure 2.**
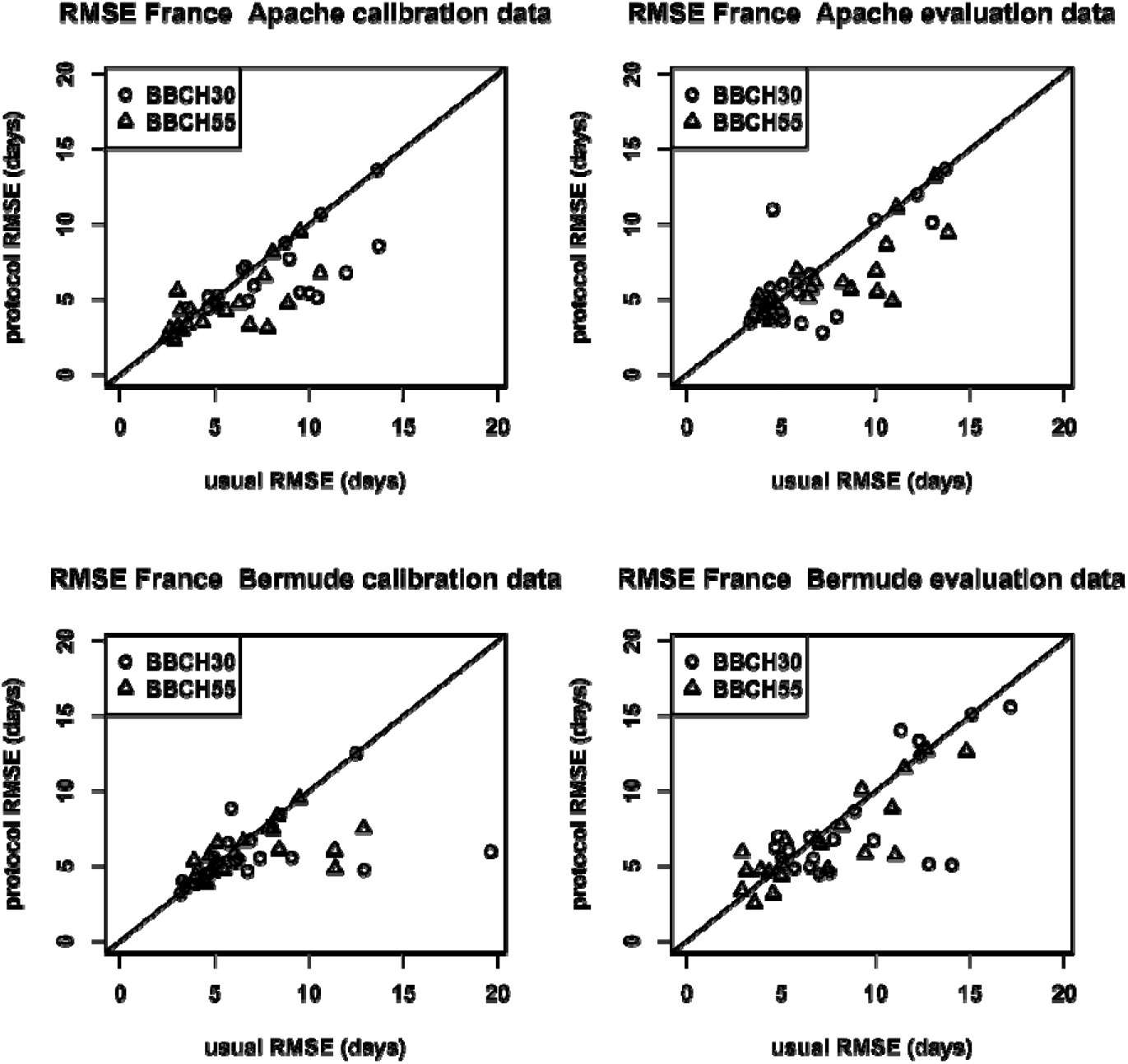
Root mean squared error (RMSE) of results following usual or protocol calibration, for each modeling team, separately for calibration and evaluation data of the French data sets.

**Table 7:** First example of documentation for protocol step Step 6, “Examine goodness-of-fit”. In this table there is one row for each observed variable with a simulated equivalent, showing mean squared error (MSE) and its decomposition into three terms. Of particular interest is the bias^2^ contribution, which should be small if there is an almost obligatory parameter corresponding to this variable. This example is for the French data set, variety Apache, modeling team M21.

|  | MSE<br>(days<sup>2</sup>) | bias<sup>2</sup><br>(days<sup>2</sup>) | SDSD<br>(days<sup>2</sup>) | LCS<br>(days<sup>2</sup>) |
| --- | --- | --- | --- | --- |
| BBCH30 | 19.64 | 0.25 | 5.93 | 13.46 |
| BBCH55 | 5.29 | 0.02 | 0.03 | 5.24 |

**Table 8:** Second example of documentation for protocol Step 6, “Examine goodness-of-fit”. In this table there is one row for each observed variable with a simulated equivalent, showing root mean squared error (RMSE) for the calibrated model and for two benchmark models. The “naïve” benchmark assumes that all environments have the same number of days to the given development stage, equal to the average of the observed days to that stage. The “onlyT” benchmark assumes a constant number of degree days to the stage in question, with a simple vernalization calculation in the case of winter wheat. This example is for the French data set, variety Apache, modeling team M21.

|  | naive | onlyT | M21 |
| --- | --- | --- | --- |
| variable | RMSE<br>(days) | RMSE<br>(days) | RMSE<br>(days) |
| BBCH30 | 12.5 | 8.4 | 3.1 |
| BBCH55 | 8.3 | 9.5 | 3.8 |

### Comparison of protocol and usual calibration

Using usual or protocol calibration led to important differences in the calibrated models. For example, the number of estimated parameters in the final model was different between protocol and usual calibration (Supplementary Figure S1, table S1). The differences between simulated values after usual and protocol calibration were small for BBCH10, for which there were no calibration data. For the other stages, the simulated values differed appreciably (Supplementary Figure S2, Table S2).

### Goodness of fit and evaluation for usual and protocol calibration

Figures 2 and 3 show RMSE using usual and protocol calibration for the French and Australian data sets respectively (results by modeling team are in Supplementary Tables S3-S8). Table 9 shows RMSE values for each data set, averaged over modeling teams, for usual and protocol calibration for the calibration and evaluation data. The protocol reduces RMSE by 10-22% compared to the usual calibration method. The *p* values for a one-sided paired *t*-test of the hypothesis that RMSE is larger for usual calibration than for protocol calibration are also shown. On average over stages other than BBCH10, all three data sets have significantly larger RMSE values with usual calibration than with protocol calibration for the calibration data (*p*<0.05). For the evaluation data, p<0.01 for the two French data sets, but *p*=0.15 for the Australian data set. The table also shows the proportion of modeling teams where RMSE is larger for the usual calibration than for the protocol calibration. Looking at the averages over stages and then averaging over data sets, 75% of models have lower RMSE for protocol calibration than for usual calibration for the calibration data, and 66% for the evaluation data. Almost all modeling teams did better than the two benchmark models for all stages, both for usual calibration and protocol calibration, with slightly better results for protocol calibration (Supplementary Tables S3-S8). Since the protocol specifically aims to reduce bias, one would expect squared bias to be a smaller fraction of MSE for protocol calibration than for usual calibration, and this is the case, both for the calibration data and the evaluation data (Supplementary Tables S9-S23).

**Figure 3.**
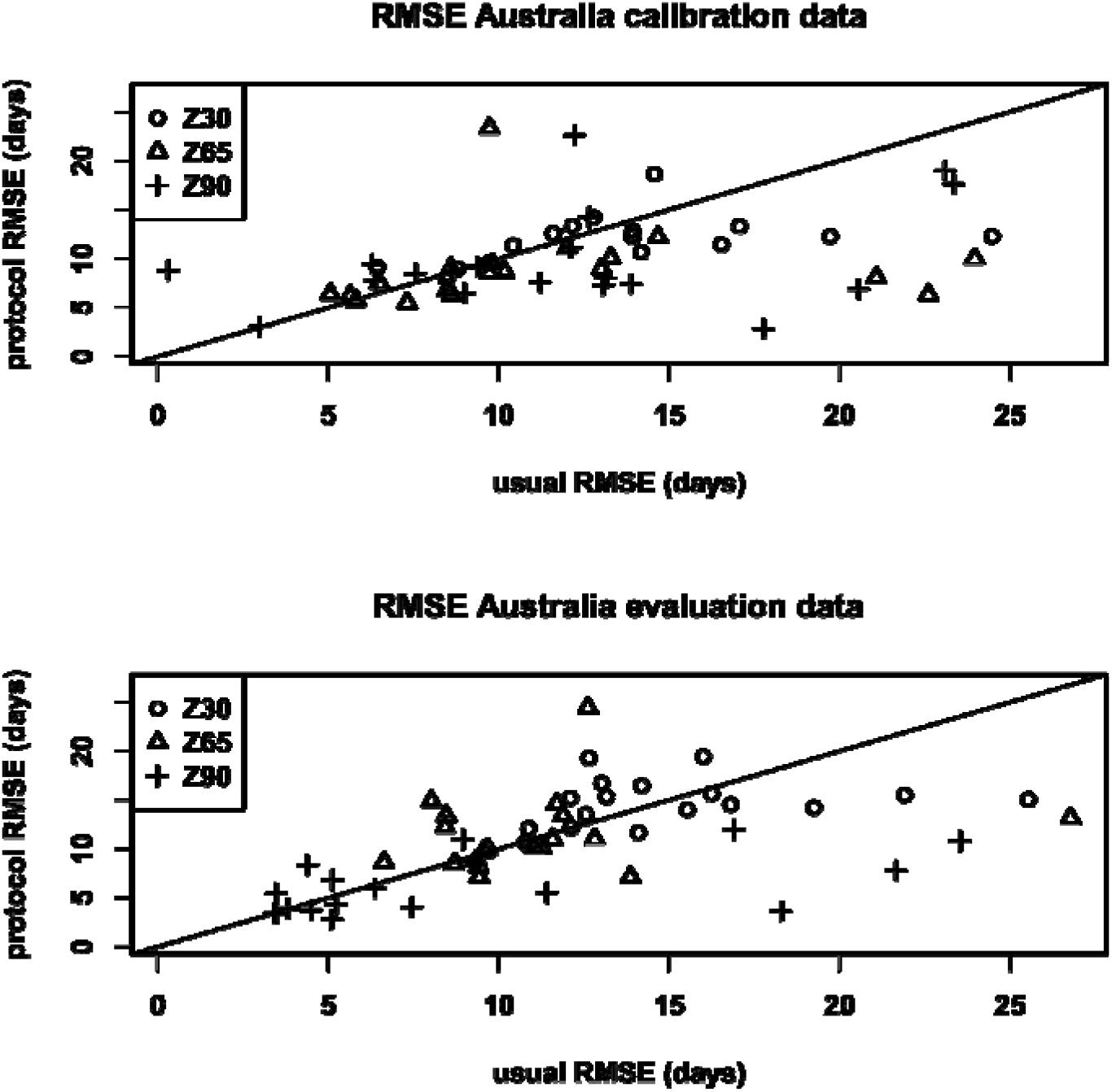
Root mean squared error (RMSE) of results following usual or protocol calibration, for each modeling team, separately for calibration and evaluation data of the Australian data set.

**Table 9:**
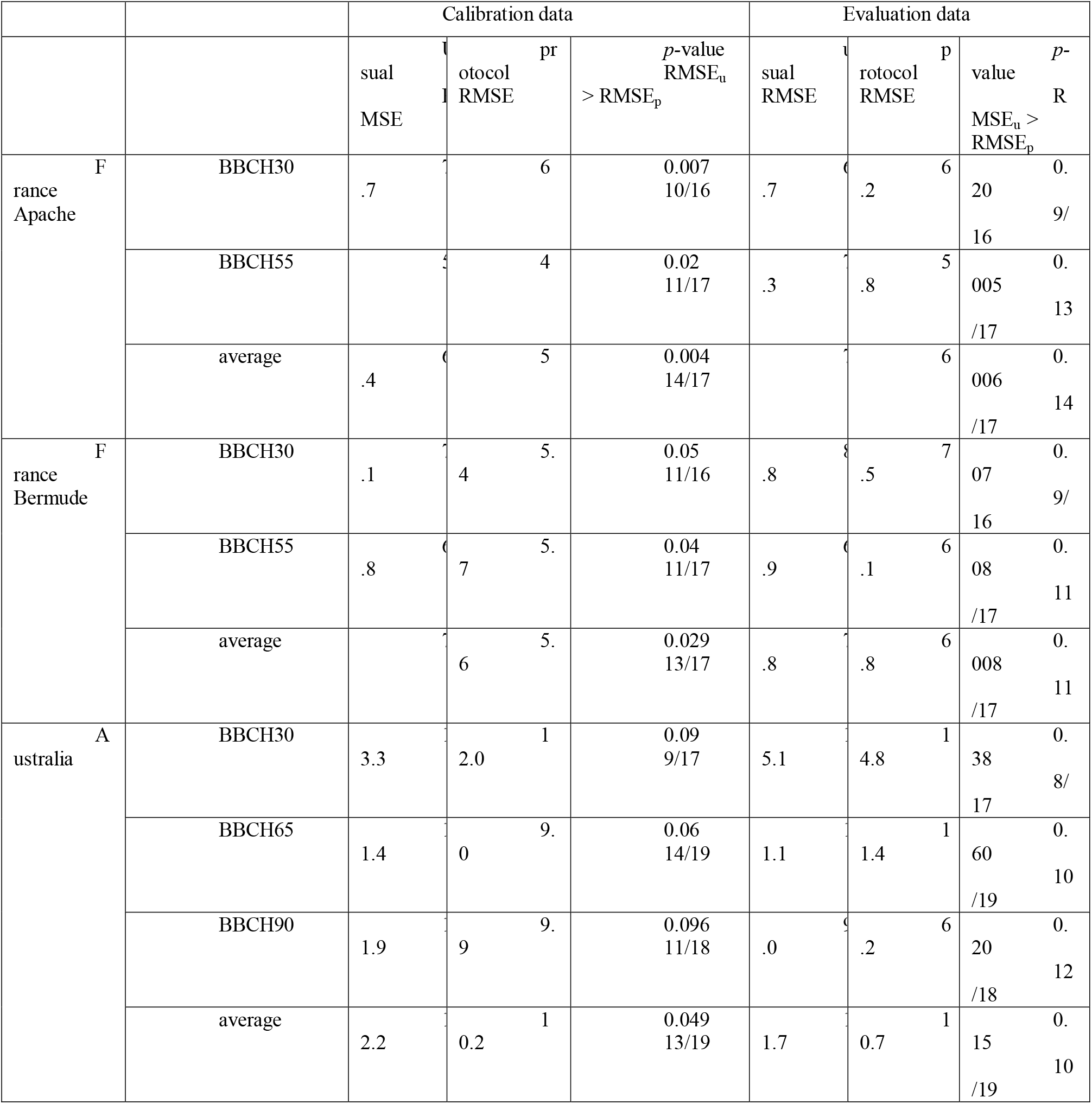
Comparison of errors for usual and protocol calibration. The table shows root mean squared error (RMSE) averaged over modeling teams for each stage and for the average over stages, separately for the calibration and evaluation data. For each of calibration and evaluation data, the first column is RMSE for simulations using the usual calibration method, the second column is RMSE using protocol calibration, and the third column is the p value of a one sided paired t test, that tests whether RMSE for usual calibration is larger than for protocol calibration. Below the p value is the fraction of modeling teams for which RMSE for usual calibration is larger than for protocol calibration.

### Comparison of usual and protocol calibration for ensemble of models

The choice of usual or protocol calibration has little effect on the predictive accuracy of the ensemble models e-mean and e-median. Averaged over development stages and over data sets, for the evaluation data, RMSE for e-median is respectively 5.7 and 5.8 days for usual and protocol calibration. The values for RMSE of e-mean are 6.1 and 6.2 days for usual and protocol calibration, respectively (Supplementary Tables S4, S6, S8).

Recently, many crop model studies have been based on ensembles of models (Jägermeyr et al. 2021). Many studies have found that the ensemble mean and median are good predictors, sometimes better than even the best individual model (Martre et al. 2015; Wallach et al. 2018; Farina et al. 2021). It has thus, become quite common to base projections of climate change impact on crop production on the ensemble median (e.g. Asseng et al., 2019). The e-mean and e-median results here, compared to the individual modeling teams, are in keeping with previous results. The e-median model is among the better predictors though not the very best, and is somewhat better than e-mean.

The variability between simulated results for different modeling teams is shown in Table 10. The standard deviation is similar for usual and protocol calibration for BBCH10, for which there are no data for calibration, but is systematically smaller for protocol calibration for the other stages. Considering the average over stages other than BBCH10 and taking the average over data sets, protocol calibration reduced the standard deviation of simulated values by 31% for the calibration data and by 22% for the evaluation data.

**Table 10:**
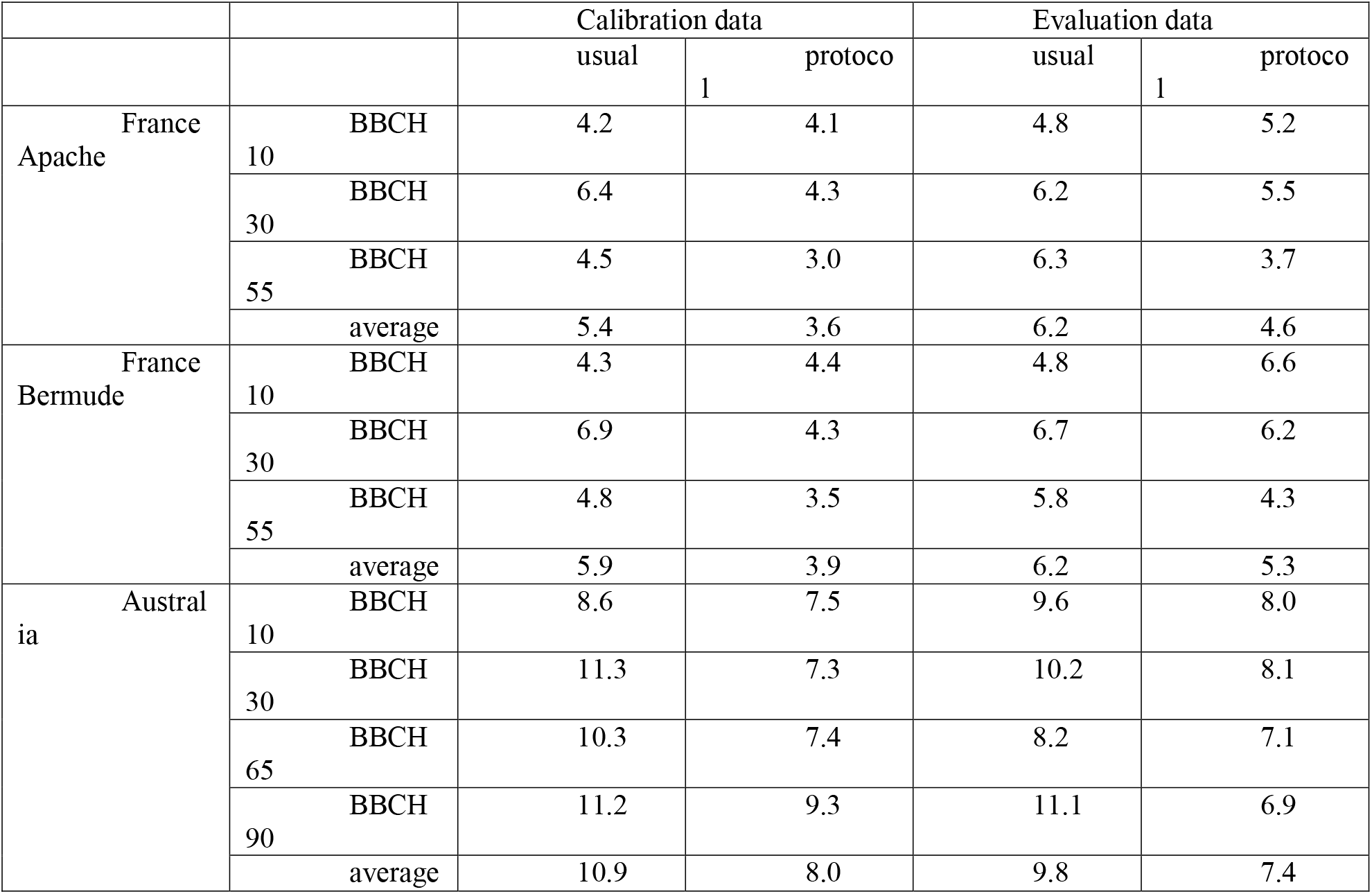
Variability of simulated values. This table shows the standard deviations of values simulated by the different modeling teams (days), for each simulated development stage and for the average over stages other than BBCH10. BBCH is not considered, since there were no observed values for BBCH10. Separate standard deviations are given for the calibration and evaluation data, and simulation using usual calibration or protocol calibration.

## 4. Conclusions

We propose an original, detailed protocol for calibration of crop phenology models and show that it can be applied by a wide range of wheat models and for data sets with different structures. While the application here is to wheat phenology models, the same protocol could undoubtedly be used more generally for phenology models of other crops.

Comparison with usual calibration practices shows that, on average over modeling teams, the protocol leads to a better fit to the calibration data and to a better fit to out-of-sample data. Application of the protocol would be advantageous not just for individual modeling studies, but also for studies based on ensembles of models, including projections of climate impact. In particular, we have shown that if all modeling teams use the protocol, between-model variability can be substantially reduced, thus reducing the uncertainty of projections.

## Author Declarations

### Code availability / Data availability

The datasets generated during and/or analyzed during the current study are not publicly available but are available from the authors on reasonable request.

### Ethics approval/declarations

Not applicable

### Consent to participate

Not applicable

### Consent for publication

Not applicable

### Authors’ contributions

Daniel Wallach: Conceptualization, methodology, project administration, writing - original draft, validation. Taru Palosuo, Henrike Mielenz, Peter Thorburn, Samuel Buis, Sabine Seidel: Conceptualization, methodology, project administration, writing, review & editing. All other authors: Simulations, model expertise, writing, review & editing.

## References

Ahuja LR, Ma L (eds) (2011) Methods of Introducing System Models into Agricultural Research. American Society of Agronomy and Soil Science Society of America, Madison, WI, USA

Asseng S, Ewert F, Rosenzweig C, et al (2013) Uncertainty in simulating wheat yields under climate change. Nat Clim Chang 3:827–832. doi: 10.1038/nclimate1916

Asseng S, Martre P, Maiorano A, et al (2019) Climate change impact and adaptation for wheat protein. Glob Chang Biol 25:155–173. doi: 10.1111/gcb.14481

Bassu S, Brisson N, Durand J-L, et al (2014) How do various maize crop models vary in their responses to climate change factors? Glob Chang Biol 20:2301–20. doi: 10.1111/gcb.12520

Brisson N, Gary C, Justes E, et al (2003) An overview of the crop model stics. Eur J Agron 18:309–332. doi: 10.1016/S1161-0301(02)00110-7

Buis S, Lecharpentier P, Vezy R, et al (2021) SticsRPacks/CroptimizR: v0.4.0. doi: 10.5281/ZENODO.5121194

Chakrabarti A, Ghosh JK (2011) AIC, BIC and Recent Advances in Model Selection. Philos Stat 583–605. doi: 10.1016/B978-0-444-51862-0.50018-6

Cleland EE, Chuine I, Menzel A, et al (2007) Shifting plant phenology in response to global change. Trends Ecol. Evol. 22:357–365

Confalonieri R, Orlando F, Paleari L, et al (2016) Uncertainty in crop model predictions: What is the role of users? Environ Model Softw 81:165–173. doi: 10.1016/j.envsoft.2016.04.009

Farina R, Sándor R, Abdalla M, et al (2021) Ensemble modelling, uncertainty and robust predictions of organic carbon in long □ term bare □ fallow soils. Glob Chang Biol 27:904–928. doi: 10.1111/gcb.15441

Fath B, Jorgensen SE (2011) Fundamentals of ecological modelling: Applications in environmental management and research. 4th edition. Elsevier, Amsterdam

Gao Y, Wallach D, Hasegawa T, et al (2021) Evaluation of crop model prediction and uncertainty using Bayesian parameter estimation and Bayesian model averaging. Agric For Meteorol 311:108686. doi: 10.1016/J.AGRFORMET.2021.108686

Jägermeyr J, Müller C, Ruane AC, et al (2021) Climate impacts on global agriculture emerge earlier in new generation of climate and crop models. Nat Food 2:873–885. doi: 10.1038/s43016-021-00400-y

Khorashadi Zadeh F, Nossent J, Woldegiorgis BT, et al (2022) A fast and effective parameterization of water quality models. Environ Model Softw 149:105331. doi: 10.1016/J.ENVSOFT.2022.105331

Kobayashi K (2004) Comments on another way of partitioning mean squared deviation proposed by Gauch et al. (2003). With reply. Agron J 96:1206–1207

Kuha J (2004) AIC and BIC: Comparisons of Assumptions and Performance. Sociol Methods Res 33:188–229. doi: 10.1177/0049124103262065

Lawes RA, Huth ND, Hochman Z (2016) Commercially available wheat cultivars are broadly adapted to location and time of sowing in Australia’s grain zone. Eur J Agron 77:38–46. doi: 10.1016/J.EJA.2016.03.009

Li T, Hasegawa T, Yin X, et al (2015) Uncertainties in predicting rice yield by current crop models under a wide range of climatic conditions. Glob Chang Biol 21:1328–41. doi: 10.1111/gcb.12758

Martre P, Wallach D, Asseng S, et al (2015) Multimodel ensembles of wheat growth: many models are better than one. Glob Chang Biol 21:911–25. doi: 10.1111/gcb.12768

McNunn G, Heaton E, Archontoulis S, et al (2019) Using a Crop Modeling Framework for Precision Cost-Benefit Analysis of Variable Seeding and Nitrogen Application Rates. Front Sustain Food Syst 3:108. doi: 10.3389/fsufs.2019.00108

Menzel A, Yuan Y, Matiu M, et al (2020) Climate change fingerprints in recent European plant phenology. Glob Chang Biol 26:2599–2612. doi: 10.1111/gcb.15000

Nelder JA, Mead R (1965) A Simplex Method for Function Minimization. Comput J 7:308–313. doi: 10.1093/comjnl/7.4.308

Piao S, Liu Q, Chen A, et al (2019) Plant phenology and global climate change: current progresses and challenges. Glob Chang Biol gcb.14619. doi: 10.1111/gcb.14619

Rafiei V, Nejadhashemi AP, Mushtaq S, et al (2022) An improved calibration technique to address high dimensionality and non-linearity in integrated groundwater and surface water models. Environ Model Softw 149:105312. doi: 10.1016/J.ENVSOFT.2022.105312

Rezaei EE, Siebert S, Hüging H, Ewert F (2018) Climate change effect on wheat phenology depends on cultivar change. Sci Rep 8:4891. doi: 10.1038/s41598-018-23101-2

Sisheber B, Marshall M, Mengistu D, Nelson A (2022) Tracking crop phenology in a highly dynamic landscape with knowledge-based Landsat–MODIS data fusion. Int J Appl Earth Obs Geoinf 106:102670. doi: 10.1016/J.JAG.2021.102670

Stuble KL, Bennion LD, Kuebbing SE (2021) Plant phenological responses to experimental warming – A synthesis. Glob Chang Biol gcb.15685. doi: 10.1111/gcb.15685

Tao F, Rötter RP, Palosuo T, et al (2018) Contribution of crop model structure, parameters and climate projections to uncertainty in climate change impact assessments. Glob Chang Biol 24:1291–1307. doi: 10.1111/gcb.14019

van Bussel LGJ, Stehfest E, Siebert S, et al (2015) Simulation of the phenological development of wheat and maize at the global scale. Glob Ecol Biogeogr 24:1018–1029. doi: 10.1111/geb.12351

Wallach D (2011) Crop model calibration: A statistical perspective. Agron J 103:1144–1151

Wallach D, Martre P, Liu B, et al (2018) Multimodel ensembles improve predictions of crop-environment-management interactions. Glob Chang Biol 24:5072–5083. doi: 10.1111/gcb.14411

Wallach D, Palosuo T, Thorburn P, et al (2021a) How well do crop modeling groups predict wheat phenology, given calibration data from the target population? Eur J Agron 124:126195. doi: https://doi.org/10.1016/j.eja.2020.126195

Wallach D, Palosuo T, Thorburn P, et al (2021b) Multi-model evaluation of phenology prediction for wheat in Australia. Agric For Meteorol 298-299:108289. doi: 10.1016/J.AGRFORMET.2020.108289

Wallach D, Palosuo T, Thorburn P, et al (2021c) The chaos in calibrating crop models: Lessons learned from a multi-model calibration exercise. Environ Model Softw 145:105206. doi: 10.1016/J.ENVSOFT.2021.105206

Wallach D, Palosuo T, Thorburn P, et al (2021d) Multi model evaluation of phenology prediction for wheat in Australia. Agric For Meteorol in press: doi: 10.1101/2020.06.06.133504

Wang PC, Shoup TE (2011) Parameter sensitivity study of the Nelder–Mead Simplex Method. Adv Eng Softw 42:529–533. doi: 10.1016/J.ADVENGSOFT.2011.04.004

Webber H, Lischeid G, Sommer M, et al (2020) No perfect storm for crop yield failure in Germany. Environ Res Lett 15:. doi: 10.1088/1748-9326/aba2a4

Zhang L, Zhang Z, Tao F, et al (2022) Adapting to climate change precisely through cultivars renewal for rice production across China: When, where, and what cultivars will be required? Agric For Meteorol 316:108856. doi: 10.1016/J.AGRFORMET.2022.108856

