## Supplementary material for "Proposal and extensive test of a calibration protocol for crop phenology models": SI

### **Supplementary information**

Running title: Phenology model calibration

Daniel Wallach^1^, Taru Palosuo^2^, Peter Thorburn^3^, Henrike Mielenz^4^, Samuel Buis^5^, Zvi Hochman^3^, Emmanuelle Gourdain^6^, Fety Andrianasolo^6^, Benjamin Dumont^7^, Roberto Ferrise^8^, Thomas Gaiser^9^, Cecile Garcia^6^, Sebastian Gayler^10^, Matthew Harrison^11^, Santosh Hiremath^12^, Heidi Horan^3^, Gerrit Hoogenboom^13,14^, Per-Erik Jansson^15^, Qi Jing^16^, Eric Justes^17^, Kurt-Christian Kersebaum^18,19^, Marie Launay^20^, Elisabet Lewan^20^, Ke Liu^11^, Fasil Mequanint^10^, Marco Moriondo^22^, Claas Nendel^18,19,23^, Gloria Padovan^8^, Budong Qian^16^, Niels Schütze^24^, Diana-Maria Seserman^18^, Vakhtang Shelia^13,14^, Amir Souissi^25^, Xenia Specka^18^, Amit Kumar Srivastava^9^, Giacomo Trombi^8^, Tobias K.D. Weber^10^, Lutz Weihermüller^26^, Thomas Wöhling^24,27^, Sabine J. Seidel ^9*^

^1^INRAE, UMR AGIR, Castanet Tolosan, France. ORCID 0000-0003-3500-8179

^2^Natural Resources Institute Finland (Luke), Helsinki, Finland

^3^CSIRO Agriculture and Food, Brisbane, Queensland, Australia

^4^Institute for Crop and Soil Science, Federal Research Centre for cultivated Plants, Julius Kühn-Institut (JKI), Braunschweig, Germany

^5^INRAE, UMR 1114 EMMAH, Avignon, France

^6^ARVALIS - Institut du végétal Paris, France

^7^Plant Sciences & TERRA Teaching and Research Centre, Gembloux Agro-Bio Tech, University of Liege, Gembloux, Belgium

^8^Department of Agriculture, Food, Environment and Forestry (DAGRI), University of Florence, Italy

^9^Institute of Crop Science and Resource Conservation, University of Bonn, Germany

^10^Institute of Soil Science and Land Evaluation, Biogeophysics, University of Hohenheim, Stuttgart, Germany

^11^Tasmanian Institute of Agriculture, University of Tasmania Launceston, Australia

^12^Aalto University School of Science, Espoo, Finland;

^13^Agricultural and Biological Engineering Department, University of Florida, Gainesville, Florida, USA

^14^Food Systems Institute, University of Florida, Gainesville, Florida, USA

^15^Royal Institute of Technology (KTH), Stockholm, Sweden

^16^Ottawa Research and Development Centre, Agriculture and Agri-Food Canada, Ottawa, Canada

^17^CIRAD, UMR SYSTEM, Montpellier, France

^18^Leibniz Centre for Agricultural Landscape Research (ZALF), Müncheberg, Germany

^19^Global Change Research Institute CAS, Brno, Czech Republic^19^INRAE, US 1116 AgroClim, Avignon, France

^20^INRAE, US 1116 AgroClim, Avignon, France

^21^Department of Soil and Environment, Swedish University of Agricultural Sciences (SLU), Uppsala, Sweden

^22^CNR-IBE, Firenze, Italy

^23^Institute of Biochemistry and Biology, University of Potsdam, Potsdam, Germany

^24^Institute of Hydrology and Meteorology, Chair of Hydrology, Technische Universität Dresden, Dresden, Germany

^25^Swift Current Research and Development Centre, Agriculture and Agri-Food Canada, Swift Current, Saskatchewan, Canada.

^26^Institute of Bio- and Geosciences - IBG-3, Agrosphere, Forschungszentrum Jülich GmbH, Jülich, Germany

^27^Lincoln Agritech Ltd., Hamilton, New Zealand

Figure 1

**Number of parameters in the final calibrated model for usual and protocol calibration, for each dataset.**

Figure 2

**Usual versus protocol simulated values for each dataset. Points outside the dotted lines have a difference of more than 10 days for the French datasets and more than 20 days for the Australian dataset.**

TableS1

Number of estimated parameters in the final model after protocol or usual calibration, averaged over modeling teams

| Data set | Protocol calibration | | | | Usual calibration |
| --- | --- | --- | --- | --- | --- |
|  | Additive parameters | Candidate parameters | Accepted candidate parameters | Total parameters | Total parameters |
| France  Apache | 1.9 | 3.6 | 0.7 | 2.6 | 3.5 |
| France  Bermude | 1.9 | 3.6 | 0.9 | 2.8 | 3.6 |
| Australia | 4.0 | 4.5 | 1.7 | 5.7 | 5.7 |

Table S2

Mean over modeling teams and environments of the absolute difference in simulated values between usual calibration and protocol calibration for the calibration data.

| French data set (Apache and Bermude) | stage | BBCH10 | BBCH30 | BBCH55 |  |
| --- | --- | --- | --- | --- | --- |
|  | Mean absolute difference (days) | 0.8 | 4.5 | 3.1 |  |
| Australian data set | stage | BBCH10 | BBCH30 | BBCH65 | BBCH90 |
|  | Mean absolute difference (days) | 1.8 | 7.9 | 7.9 | 8.5 |

Table S3

**RMSE for usual and protocol calibration, for Apache calibration dataset. Models are sorted in order of increasing RMSE for the Apache evaluation data averaged over stages (last column of Table 3).**

|  | Apache calibration data | | | | | |
| --- | --- | --- | --- | --- | --- | --- |
|  | usual | | | protocol | | |
| model | BBCH30 | BBCH55 | average | BBCH30 | BBCH55 | average |
| M7 | 9.5 | 3.6 | 6.6 | 5.5 | 3.3 | 4.4 |
| M8 | 10.4 | 3.1 | 6.8 | 5.1 | 3.2 | 4.1 |
| e-median | 5.1 | 2.6 | 3.9 | 4.7 | 3 | 3.9 |
| e-mean | 5.1 | 3.2 | 4.2 | 4.5 | 3 | 3.8 |
| M20 | 6.8 | 3.2 | 5 | 4.9 | 4.3 | 4.6 |
| M12 | 10 | 7.8 | 8.9 | 5.4 | 3.1 | 4.3 |
| M16 | 4.7 | 6.9 | 5.8 | 5.2 | 3.3 | 4.2 |
| M13 | 7.1 | 4.4 | 5.7 | 5.9 | 3.5 | 4.7 |
| M17 | 5 | 2.5 | 3.7 | 4.8 | 2.4 | 3.6 |
| M15 | 5.2 | 2.8 | 4 | 5.2 | 2.6 | 3.9 |
| M25 | 6.5 | 3 | 4.8 | 7 | 5.6 | 6.3 |
| M26 | 3.6 | 3.7 | 3.7 | 4.4 | 4.3 | 4.3 |
| M21 | 4.6 | 2.9 | 3.7 | 4.4 | 2.3 | 3.4 |
| M24 | 9 | 5.6 | 7.3 | 7.7 | 4.2 | 6 |
| M9 | NA | 6.3 | 6.3 | NA | 4.7 | 4.7 |
| M2 | 6.6 | 3 | 4.8 | 7.2 | 3.1 | 5.2 |
| M30 | 12 | 8.9 | 10.4 | 6.7 | 4.7 | 5.7 |
| M1 | 8.8 | 7.6 | 8.2 | 8.8 | 6.5 | 7.6 |
| M6 | 13.7 | 10.6 | 12.1 | 8.6 | 6.7 | 7.6 |
| onlyT | 10.7 | 8.1 | 9.4 | 10.7 | 8.1 | 9.4 |
| naive | 13.6 | 9.5 | 11.6 | 13.6 | 9.5 | 11.6 |

**Table S4**

**RMSE for usual and protocol calibration, for Apache evaluation dataset. Models are sorted in order of increasing RMSE averaged over stages. (lasr column).**

|  | Apache evaluation data | | | | | |
| --- | --- | --- | --- | --- | --- | --- |
|  | usual | | | protocol | | |
| model | BBCH30 | BBCH55 | average | BBCH30 | BBCH55 | average |
| M7 | 7.2 | 3.8 | 5.5 | 2.8 | 5.2 | 4 |
| M8 | 6.1 | 3.8 | 4.9 | 3.4 | 4.8 | 4.1 |
| e-median | 3.3 | 4.4 | 3.9 | 3.4 | 4.8 | 4.1 |
| e-mean | 3.5 | 4.7 | 4.1 | 3.9 | 4.6 | 4.3 |
| M20 | 7.9 | 6.4 | 7.2 | 3.9 | 5.1 | 4.5 |
| M12 | 4.1 | 10.1 | 7.1 | 4.1 | 5.5 | 4.8 |
| M16 | 5 | 8.7 | 6.9 | 4.1 | 5.7 | 4.9 |
| M13 | 4.6 | 6.8 | 5.7 | 3.7 | 6.1 | 4.9 |
| M17 | 5.1 | 4.1 | 4.6 | 6 | 3.9 | 4.9 |
| M15 | 5.8 | 4.8 | 5.3 | 5.4 | 4.6 | 5 |
| M25 | 5.1 | 5.8 | 5.5 | 3.6 | 6.9 | 5.3 |
| M26 | 6.5 | 4.5 | 5.5 | 6.6 | 5 | 5.8 |
| M21 | 5.8 | 6.6 | 6.2 | 6 | 5.7 | 5.8 |
| M24 | 4.4 | 8.3 | 6.3 | 5.7 | 6.1 | 5.9 |
| M9 | NA | 10 | 10 | NA | 6.9 | 6.9 |
| M2 | 4.5 | 4.4 | 4.5 | 11 | 3.6 | 7.3 |
| M30 | 12.2 | 10.9 | 11.5 | 12 | 4.9 | 8.4 |
| M1 | 10 | 10.6 | 10.3 | 10.3 | 8.6 | 9.5 |
| M6 | 13 | 13.8 | 13.4 | 10.1 | 9.4 | 9.8 |
| onlyT | 13.3 | 11.1 | 12.2 | 13.3 | 11.1 | 12.2 |
| naive | 13.7 | 13.1 | 13.4 | 13.7 | 13.1 | 13.4 |

Table S5

**RMSE for usual and protocol calibration, for Bermude calibration dataset. Models are sorted in order of increasing RMSE for the Bermude evaluation data averaged over stages (last column of Table 5).**

|  | Bermude calibration data | | | | | |
| --- | --- | --- | --- | --- | --- | --- |
|  | usual | | | protocol | | |
| model | BBCH30 | BBCH55 | average | BBCH30 | BBCH55 | average |
| M15 | 3.2 | 3.8 | 3.5 | 3.1 | 3.8 | 3.5 |
| M2 | 4.5 | 4.9 | 4.7 | 4.5 | 4.6 | 4.5 |
| M17 | 6.3 | 4.4 | 5.4 | 5.4 | 4 | 4.7 |
| e-median | 3.3 | 5 | 4.1 | 4 | 5 | 4.5 |
| M8 | 7.4 | 4 | 5.7 | 5.5 | 4.4 | 4.9 |
| M21 | 4 | 4.6 | 4.3 | 4.1 | 3.9 | 4 |
| M12 | 12.9 | 11.4 | 12.2 | 4.7 | 4.8 | 4.8 |
| e-mean | 3.4 | 5.5 | 4.5 | 3.6 | 4.8 | 4.2 |
| M7 | 6.8 | 3.9 | 5.3 | 4.7 | 5.4 | 5 |
| M25 | 4.7 | 4.7 | 4.7 | 4.4 | 5.6 | 5 |
| M16 | 5 | 11.4 | 8.2 | 5.6 | 6 | 5.8 |
| M13 | 6.1 | 5.2 | 5.6 | 5.3 | 6.5 | 5.9 |
| M26 | 5.7 | 6.5 | 6.1 | 6.5 | 6.6 | 6.6 |
| M20 | 5.2 | 6.1 | 5.6 | 5.2 | 5.7 | 5.5 |
| M24 | 9.1 | 8.1 | 8.6 | 5.6 | 7.4 | 6.5 |
| M9 | NA | 8.4 | 8.4 | NA | 6.1 | 6.1 |
| M30 | 5.9 | 7.8 | 6.9 | 8.8 | 7.5 | 8.2 |
| M1 | 6.9 | 8.1 | 7.5 | 6.7 | 7.5 | 7.1 |
| naive | 12.5 | 8.3 | 10.4 | 12.5 | 8.3 | 10.4 |
| onlyT | 8.4 | 9.5 | 9 | 8.4 | 9.5 | 9 |
| M6 | 19.6 | 12.9 | 16.3 | 5.9 | 7.5 | 6.7 |

Table S6

**RMSE for usual and protocol calibration, for Bermude evaluation dataset. Models are sorted in order of increasing RMSE averaged over stages (last column).**

|  | Bermude evaluation data | | | | | |
| --- | --- | --- | --- | --- | --- | --- |
|  | usual | | | protocol | | |
| model | BBCH30 | BBCH55 | average | BBCH30 | BBCH55 | average |
| M15 | 5 | 2.9 | 4 | 5.5 | 3.4 | 4.4 |
| M2 | 7 | 4.4 | 5.7 | 4.5 | 4.6 | 4.5 |
| M17 | 4.8 | 3.6 | 4.2 | 7 | 2.5 | 4.7 |
| e-median | 6.5 | 3.9 | 5.2 | 4.9 | 4.7 | 4.8 |
| M8 | 12.8 | 3.2 | 8 | 5.1 | 4.7 | 4.9 |
| M21 | 6.5 | 4.6 | 5.5 | 6.9 | 3.1 | 5 |
| M12 | 7.6 | 11 | 9.3 | 4.6 | 5.7 | 5.1 |
| e-mean | 6.7 | 5 | 5.9 | 5.5 | 5.1 | 5.3 |
| M7 | 14.1 | 2.9 | 8.5 | 5 | 5.9 | 5.5 |
| M25 | 5.7 | 7.1 | 6.4 | 4.8 | 6.5 | 5.7 |
| M16 | 7.8 | 9.4 | 8.6 | 6.8 | 5.8 | 6.3 |
| M13 | 5.4 | 5.2 | 5.3 | 6 | 6.7 | 6.4 |
| M26 | 8.9 | 5 | 7 | 8.7 | 4.4 | 6.5 |
| M20 | 9.9 | 6.9 | 8.4 | 6.7 | 6.8 | 6.7 |
| M24 | 4.7 | 8.2 | 6.4 | 6.2 | 7.7 | 7 |
| M9 | NA | 10.9 | 10.9 | NA | 8.9 | 8.9 |
| M30 | 11.3 | 7.5 | 9.4 | 14 | 4.8 | 9.4 |
| M1 | 12.3 | 9.3 | 10.8 | 13.3 | 10.1 | 11.7 |
| naive | 12.3 | 11.5 | 11.9 | 12.3 | 11.5 | 11.9 |
| onlyT | 15.1 | 12.7 | 13.9 | 15.1 | 12.7 | 13.9 |
| M6 | 17.2 | 14.8 | 16 | 15.6 | 12.6 | 14.1 |

Table S7

**RMSE for usual and protocol calibration, for Australian calibration dataset. Models are sorted in order of increasing RMSE for the Australian evaluation data averaged over stages (last column of Table 7).**

|  | Australian calibration data | | | | | | | |
| --- | --- | --- | --- | --- | --- | --- | --- | --- |
|  | Usual calibration | | | | Protocol calibration | | | |
| model | Z30 | Z65 | Z90 | average | Z30 | Z65 | Z90 | average |
| M9 | NA | 5.6 | 11.2 | 8.4 | NA | 6.2 | 7.6 | 6.9 |
| M17 | 8.5 | 8.6 | 17.8 | 11.6 | 8 | 9.1 | 2.9 | 6.6 |
| M13 | 14.2 | 10.2 | 13.1 | 12.5 | 10.7 | 8.7 | 7.3 | 8.9 |
| e-median | 9.3 | 5.7 | 7.2 | 7.4 | 9.9 | 6.2 | 7.3 | 7.8 |
| e-mean | 9.8 | 5.8 | 8.4 | 8 | 10 | 6.1 | 7.8 | 8 |
| M20 | NA | 9.7 | 12.1 | 10.9 | NA | 8.6 | 11.1 | 9.9 |
| M15 | 12.2 | 13.3 | 3 | 9.5 | 13.4 | 10.1 | 3.1 | 8.8 |
| M2 | 11.6 | 5.8 | 9.4 | 8.9 | 12.6 | 5.7 | 9.2 | 9.2 |
| M24 | 9.8 | 9.8 | 12.6 | 10.7 | 9.7 | 9.5 | 14.2 | 11.1 |
| M7 | 19.7 | 24 | 13.1 | 18.9 | 12.3 | 10 | 8 | 10.1 |
| M30 | 6.4 | 12 | 0.3 | 6.3 | 9.2 | 11.1 | 8.8 | 9.7 |
| M21 | 8.8 | 7.3 | 6.4 | 7.5 | 9 | 5.5 | 7.8 | 7.4 |
| M12 | 13.9 | 8.5 | 9 | 10.5 | 12.8 | 6.8 | 6.5 | 8.7 |
| M28 | 24.5 | 22.6 | 20.6 | 22.5 | 12.3 | 6.3 | 6.9 | 8.5 |
| M25 | 12 | 5.1 | 7.6 | 8.2 | 12.3 | 6.4 | 8.5 | 9.1 |
| M8 | 16.5 | 21.1 | 13.9 | 17.2 | 11.5 | 8.1 | 7.4 | 9 |
| M26 | 14.6 | 8.6 | 6.3 | 9.8 | 18.6 | 6.4 | 9.4 | 11.5 |
| M16 | 13.9 | 14.7 | 23.3 | 17.3 | 12.3 | 12.2 | 17.6 | 14 |
| M1 | 17.1 | 6.5 | 23.1 | 15.5 | 13.4 | 7.5 | 19 | 13.3 |
| naive | 14.9 | 24.3 | 22.8 | 20.6 | 14.9 | 24.3 | 22.8 | 20.6 |
| M29 | 12.8 | 13 | NA | 12.9 | 14.2 | 9 | NA | 11.6 |
| onlyT | 13 | 15.4 | 25.8 | 18.1 | 13 | 15.4 | 25.8 | 18.1 |
| M6 | 10.4 | 9.7 | 12.2 | 10.8 | 11.4 | 23.3 | 22.6 | 19.1 |

Table S8

**RMSE for usual and protocol calibration, for Australian evaluation dataset. Models are sorted in order of increasing RMSE for the evaluation data averaged over stages (last column).**

|  | Australian evaluation data | | | | | | | |
| --- | --- | --- | --- | --- | --- | --- | --- | --- |
|  | Usual calibration | | | | Protocol calibration | | | |
| model | Z30 | Z65 | Z90 | average | Z30 | Z65 | Z90 | average |
| M9 | NA | 11 | 3.5 | 7.2 | NA | 10.2 | 5.5 | 7.8 |
| M17 | 10.9 | 9.4 | 11.4 | 10.6 | 12.1 | 7.2 | 5.5 | 8.3 |
| M13 | 14.1 | 9.6 | 3.4 | 9 | 11.7 | 9.7 | 3.4 | 8.3 |
| e-median | 11.9 | 8.5 | 3.6 | 8 | 12.2 | 9.7 | 3.3 | 8.4 |
| e-mean | 11.4 | 8.4 | 3.2 | 7.7 | 12.6 | 9.6 | 3.3 | 8.5 |
| M20 | NA | 13.9 | 8.9 | 11.4 | NA | 7.2 | 11 | 9.1 |
| M15 | 12.5 | 6.6 | 6.4 | 8.5 | 13.6 | 8.7 | 6 | 9.4 |
| M2 | 14.2 | 8.7 | 3.5 | 8.8 | 16.5 | 8.5 | 3.4 | 9.4 |
| M24 | 12.1 | 9.4 | 4.4 | 8.6 | 12.2 | 8.2 | 8.3 | 9.6 |
| M7 | 19.2 | 11.6 | 3.8 | 11.5 | 14.2 | 11.1 | 3.9 | 9.7 |
| M30 | 10.8 | 8 | 4.5 | 7.8 | 10.7 | 14.9 | 3.7 | 9.8 |
| M21 | 12.1 | 8.4 | 5.1 | 8.5 | 15.2 | 12.2 | 2.8 | 10.1 |
| M12 | 15.5 | 9.4 | 9.3 | 11.4 | 14 | 8.8 | 8.7 | 10.5 |
| M28 | 25.5 | 26.8 | 18.3 | 23.5 | 15.1 | 13.2 | 3.7 | 10.6 |
| M25 | 13.2 | 8.5 | 5.2 | 8.9 | 15.3 | 13.4 | 4.3 | 11 |
| M8 | 16 | 11.2 | 7.5 | 11.6 | 19.4 | 10.2 | 4 | 11.2 |
| M26 | 13 | 12.8 | 5.1 | 10.3 | 16.7 | 11.1 | 6.8 | 11.5 |
| M16 | 16.2 | 11.9 | 21.7 | 16.6 | 15.6 | 13.4 | 7.8 | 12.3 |
| M1 | 21.9 | 11.7 | 16.9 | 16.8 | 15.5 | 14.6 | 12 | 14 |
| naive | 16.8 | 17.3 | 9.9 | 14.6 | 16.8 | 17.3 | 9.9 | 14.6 |
| M29 | 12.6 | 9.7 | NA | 11.2 | 19.3 | 10 | NA | 14.7 |
| onlyT | 18.3 | 17.4 | 8.9 | 14.8 | 18.3 | 17.4 | 8.9 | 14.8 |
| M6 | 16.8 | 12.6 | 23.5 | 17.6 | 14.5 | 24.4 | 10.8 | 16.6 |

Table S9

**Decomposition of MSE into contributions from bias², SDSD, and LCS for Apache calibration data BBCH30. Models are sorted in order of increasing RMSE for the Apache evaluation data averaged over stages (last column of Table 3). Due to rounding errors, the sum of fractions from bias² SDSD and LCS may not be exactly 1.00.**

|  | Apache calibration data ,BBCH30 | | | | | | | | | |
| --- | --- | --- | --- | --- | --- | --- | --- | --- | --- | --- |
|  | Usual calibration | | | | | Protocol calibration | | | | |
| model | MSE | bias² | fraction  from bias² | fraction  from SDSD | fraction  from LCS | MSE | bias² | fraction  from bias² | fraction  from SDSD | fraction  from LCS |
| M7 | 91 | 5.22 | 0.06 | 0.01 | 0.94 | 30 | 0.08 | 0 | 0.61 | 0.39 |
| M8 | 108.86 | 0.51 | 0 | 0 | 0.99 | 26.14 | 0.33 | 0.01 | 0.52 | 0.47 |
| e-median | 25.62 | 2.36 | 0.09 | 0.3 | 0.61 | 22.25 | 0.13 | 0.01 | 0.43 | 0.56 |
| e-mean | 26.12 | 0.71 | 0.03 | 0.18 | 0.8 | 20.7 | 0.01 | 0 | 0.38 | 0.62 |
| M20 | 46.07 | 1.84 | 0.04 | 0.14 | 0.82 | 23.93 | 2.25 | 0.09 | 0.34 | 0.57 |
| M12 | 100.36 | 54.13 | 0.54 | 0.24 | 0.22 | 29.43 | 0.73 | 0.02 | 0.37 | 0.61 |
| M16 | 21.71 | 1 | 0.05 | 0 | 0.95 | 26.64 | 0.41 | 0.02 | 0.15 | 0.83 |
| M13 | 50.14 | 0.51 | 0.01 | 0.53 | 0.46 | 34.86 | 0.08 | 0 | 0.4 | 0.59 |
| M17 | 24.71 | 0.51 | 0.02 | 0.26 | 0.72 | 23.29 | 1.65 | 0.07 | 0.21 | 0.72 |
| M15 | 26.79 | 0.05 | 0 | 0.23 | 0.77 | 27.43 | 0.51 | 0.02 | 0.24 | 0.74 |
| M25 | 41.86 | 4 | 0.1 | 0.27 | 0.64 | 48.64 | 13.27 | 0.27 | 0.23 | 0.5 |
| M26 | 12.86 | 0.18 | 0.01 | 0.07 | 0.92 | 19.14 | 0.02 | 0 | 0.02 | 0.97 |
| M21 | 21.43 | 0.33 | 0.02 | 0.31 | 0.68 | 19.64 | 0.25 | 0.01 | 0.3 | 0.69 |
| M24 | 80.29 | 20.9 | 0.26 | 0.33 | 0.41 | 59.21 | 0.41 | 0.01 | 0.38 | 0.61 |
| M9 | NA | NA | NA | NA | NA | NA | NA | NA | NA | NA |
| M2 | 43.64 | 6.25 | 0.14 | 0.22 | 0.63 | 51.57 | 22.22 | 0.43 | 0.07 | 0.5 |
| M30 | 143.57 | 17.16 | 0.12 | 0.27 | 0.61 | 45.5 | 31.84 | 0.7 | 0 | 0.3 |
| M1 | 76.86 | 0.08 | 0 | 0 | 1 | 76.86 | 0.08 | 0 | 0 | 1 |
| M6 | 187.36 | 2.25 | 0.01 | 0.28 | 0.71 | 73.29 | 5.22 | 0.07 | 0 | 0.92 |
| onlyT | 113.43 | 2.04 | 0.02 | 0.15 | 0.83 | 113.43 | 2.04 | 0.02 | 0.15 | 0.83 |
| naive | 185.49 | 0 | 0 | 1 | NA | 185.49 | 0 | 0 | 1 | NA |

Table S10

**Decomposition of MSE into contributions from bias², SDSD, and LCS for Apache calibration data BBCH55. Models are sorted in order of increasing RMSE for the Apache evaluation data averaged over stages (last column of Table 3). Due to rounding errors, the sum of fractions from bias², SDSD and LCS may not be exactly 1.00.**

|  | Apache calibration data BBCH55 | | | | | | | | | |
| --- | --- | --- | --- | --- | --- | --- | --- | --- | --- | --- |
|  | Usual calibration | | | | | Protocol calibration | | | | |
| model | MSE | bias² | fraction  from bias² | fraction  from SDSD | fraction  from LCS | _MSE | bias² | fraction  from bias² | fraction  from SDSD | fraction  from LCS |
| M7 | 12.71 | 1.65 | 0.13 | 0.17 | 0.7 | 11.07 | 0.05 | 0 | 0.03 | 0.96 |
| M8 | 9.5 | 0.86 | 0.09 | 0.1 | 0.81 | 9.93 | 0.41 | 0.04 | 0.01 | 0.95 |
| e-median | 7 | 0.73 | 0.1 | 0 | 0.89 | 8.93 | 0.05 | 0.01 | 0.01 | 0.99 |
| e-mean | 10.54 | 0.8 | 0.08 | 0 | 0.92 | 8.86 | 0.25 | 0.03 | 0 | 0.97 |
| M20 | 10.14 | 1 | 0.1 | 0.02 | 0.88 | 18.14 | 4 | 0.22 | 0 | 0.78 |
| M12 | 60.86 | 49 | 0.81 | 0.01 | 0.19 | 9.64 | 0.05 | 0 | 0 | 0.99 |
| M16 | 47.07 | 16.58 | 0.35 | 0.1 | 0.54 | 10.71 | 2.04 | 0.19 | 0 | 0.81 |
| M13 | 19 | 9.88 | 0.52 | 0.03 | 0.45 | 12 | 0 | 0 | 0.01 | 0.99 |
| M17 | 6.07 | 0.05 | 0.01 | 0.01 | 0.98 | 5.93 | 0.62 | 0.1 | 0 | 0.9 |
| M15 | 8 | 0.73 | 0.09 | 0.07 | 0.83 | 6.79 | 0.62 | 0.09 | 0.08 | 0.83 |
| M25 | 9.21 | 0.86 | 0.09 | 0 | 0.9 | 31 | 16 | 0.52 | 0.05 | 0.44 |
| M26 | 13.86 | 0.02 | 0 | 0.14 | 0.86 | 18.14 | 1 | 0.06 | 0.32 | 0.62 |
| M21 | 8.21 | 0.05 | 0.01 | 0.01 | 0.99 | 5.29 | 0.02 | 0 | 0.01 | 0.99 |
| M24 | 31.43 | 13.8 | 0.44 | 0.04 | 0.52 | 17.93 | 1.15 | 0.06 | 0.03 | 0.91 |
| M9 | 39.64 | 0.25 | 0.01 | 0.02 | 0.97 | 22.29 | 0.02 | 0 | 0 | 1 |
| M2 | 9.21 | 0.25 | 0.03 | 0.02 | 0.95 | 9.86 | 0.51 | 0.05 | 0.15 | 0.8 |
| M30 | 79 | 5.9 | 0.07 | 0.27 | 0.66 | 22.43 | 0.08 | 0 | 0.2 | 0.8 |
| M1 | 58.36 | 18.98 | 0.33 | 0 | 0.67 | 42.43 | 1.65 | 0.04 | 0.01 | 0.95 |
| M6 | 112.21 | 11.27 | 0.1 | 0.36 | 0.54 | 45.14 | 0.18 | 0 | 0 | 0.99 |
| onlyT | 65.14 | 0.02 | 0 | 0.19 | 0.81 | 65.14 | 0.02 | 0 | 0.19 | 0.81 |
| naive | 90.64 | 0 | 0 | 1 | NA | 90.64 | 0 | 0 | 1 | NA |

Table S11

**Decomposition of MSE into contributions from bias², SDSD, and LCS for Bermude calibration data BBCH30. Models are sorted in order of increasing RMSE for the Bermude evaluation data averaged over stages (last column of Table 5). Due to rounding errors, the sum of fractions from bias², SDSD and LCS may not be exactly 1.00.**

|  | Bermude calibration data BBCH30 | | | | | | | | | |
| --- | --- | --- | --- | --- | --- | --- | --- | --- | --- | --- |
|  | Usual calibration | | | | | Protocol calibration | | | | |
| model | MSE | bias² | fraction  from bias² | fraction  from SDSD | fraction  from LCS | _MSE | bias² | fraction  from bias² | fraction  from SDSD | fraction  from LCS |
| M15 | 10.14 | 0.02 | 0 | 0.36 | 0.64 | 9.86 | 0.33 | 0.03 | 0.35 | 0.62 |
| M2 | 20.64 | 3.19 | 0.15 | 0.1 | 0.75 | 20.07 | 2.7 | 0.13 | 0.26 | 0.61 |
| M17 | 39.64 | 3.19 | 0.08 | 0.04 | 0.88 | 29.5 | 6.25 | 0.21 | 0.02 | 0.76 |
| e-median | 10.73 | 1.75 | 0.16 | 0.14 | 0.7 | 15.84 | 0.29 | 0.02 | 0.42 | 0.56 |
| M8 | 54.86 | 0.73 | 0.01 | 0.07 | 0.91 | 30.5 | 0.25 | 0.01 | 0.51 | 0.49 |
| M21 | 15.86 | 0.51 | 0.03 | 0.09 | 0.87 | 16.86 | 0.73 | 0.04 | 0.09 | 0.86 |
| M12 | 167.21 | 116.33 | 0.7 | 0.09 | 0.22 | 22.43 | 2.04 | 0.09 | 0.67 | 0.24 |
| e-mean | 11.81 | 2.57 | 0.22 | 0.14 | 0.64 | 12.61 | 0.13 | 0.01 | 0.3 | 0.69 |
| M7 | 45.57 | 0.02 | 0 | 0.15 | 0.85 | 21.64 | 0.01 | 0 | 0.37 | 0.63 |
| M25 | 22.14 | 0.18 | 0.01 | 0.38 | 0.61 | 19.43 | 0.02 | 0 | 0.38 | 0.62 |
| M16 | 24.64 | 8.58 | 0.35 | 0.03 | 0.62 | 31 | 18.37 | 0.59 | 0.04 | 0.36 |
| M13 | 37.64 | 0.62 | 0.02 | 0.42 | 0.57 | 27.93 | 4.9 | 0.18 | 0.3 | 0.52 |
| M26 | 32.71 | 0 | 0 | 0.01 | 0.99 | 42.5 | 0.05 | 0 | 0.01 | 0.99 |
| M20 | 26.71 | 2.04 | 0.08 | 0.08 | 0.84 | 26.79 | 1.84 | 0.07 | 0 | 0.93 |
| M24 | 82.5 | 54.13 | 0.66 | 0.19 | 0.15 | 30.86 | 0.02 | 0 | 0.44 | 0.56 |
| M9 | NA | NA | NA | NA | NA | NA | NA | NA | NA | NA |
| M30 | 34.43 | 1 | 0.03 | 0 | 0.97 | 78.14 | 59.51 | 0.76 | 0.01 | 0.23 |
| M1 | 47.86 | 0.18 | 0 | 0.02 | 0.98 | 45.07 | 2.25 | 0.05 | 0.01 | 0.94 |
| naive | 155.49 | 0 | 0 | 1 | NA | 155.49 | 0 | 0 | 1 | NA |
| onlyT | 71.29 | 1 | 0.01 | 0.38 | 0.6 | 71.29 | 1 | 0.01 | 0.38 | 0.6 |
| M6 | 385.21 | 1.47 | 0 | 0.46 | 0.53 | 35.29 | 2.94 | 0.08 | 0.06 | 0.86 |

Table S12

**Decomposition of MSE into contributions from bias², SDSD, and LCS for Bermude calibration data BBCH55. Models are sorted in order of increasing RMSE for the Bermude evaluation data averaged over stages (last column of Table 5). Due to rounding errors, the sum of fractions from bias², SDSD and LCS may not be exactly 1.00.**

|  | Bermude calibration data BBCH55 | | | | | | | | | |
| --- | --- | --- | --- | --- | --- | --- | --- | --- | --- | --- |
|  | Usual calibration | | | | | Protocol calibration | | | | |
| model | MSE | bias² | fraction  from bias² | fraction  from SDSD | fraction  from LCS | _MSE | bias² | fraction  from bias² | fraction  from SDSD | fraction  from LCS |
| M15 | 14.14 | 0 | 0 | 0.07 | 0.93 | 14.29 | 0 | 0 | 0.1 | 0.9 |
| M2 | 23.71 | 0.18 | 0.01 | 0.13 | 0.87 | 21.14 | 1 | 0.05 | 0.05 | 0.91 |
| M17 | 19.5 | 1.47 | 0.08 | 0.05 | 0.88 | 16.29 | 2.94 | 0.18 | 0.03 | 0.79 |
| e-median | 24.8 | 2.36 | 0.1 | 0.01 | 0.9 | 25.36 | 0.05 | 0 | 0.02 | 0.98 |
| M8 | 16 | 1 | 0.06 | 0 | 0.94 | 19 | 0.33 | 0.02 | 0 | 0.98 |
| M21 | 20.79 | 0.05 | 0 | 0.05 | 0.94 | 14.86 | 0.18 | 0.01 | 0.03 | 0.95 |
| M12 | 129.93 | 110.25 | 0.85 | 0 | 0.15 | 23 | 0.18 | 0.01 | 0 | 0.99 |
| e-mean | 30.09 | 4.98 | 0.17 | 0.03 | 0.8 | 22.74 | 0.06 | 0 | 0.01 | 0.99 |
| M7 | 15.29 | 0.51 | 0.03 | 0.03 | 0.93 | 28.64 | 0.13 | 0 | 0.01 | 0.99 |
| M25 | 22.29 | 1.31 | 0.06 | 0 | 0.94 | 31.57 | 1 | 0.03 | 0.11 | 0.86 |
| M16 | 129.21 | 67.47 | 0.52 | 0.09 | 0.39 | 36.43 | 12.76 | 0.35 | 0.02 | 0.63 |
| M13 | 26.57 | 6.61 | 0.25 | 0.02 | 0.73 | 42.71 | 13.8 | 0.32 | 0.01 | 0.67 |
| M26 | 42.5 | 0.01 | 0 | 0.13 | 0.87 | 43.5 | 0.25 | 0.01 | 0.3 | 0.7 |
| M20 | 36.64 | 8.58 | 0.23 | 0.07 | 0.7 | 32.93 | 1.47 | 0.04 | 0.02 | 0.94 |
| M24 | 65.57 | 36 | 0.55 | 0 | 0.45 | 54.64 | 27.19 | 0.5 | 0 | 0.5 |
| M9 | 70.79 | 7.76 | 0.11 | 0.04 | 0.85 | 37.36 | 0.05 | 0 | 0.01 | 0.99 |
| M30 | 61.57 | 12.76 | 0.21 | 0.13 | 0.66 | 56.36 | 4.29 | 0.08 | 0.14 | 0.79 |
| M1 | 65.29 | 3.45 | 0.05 | 0.03 | 0.92 | 56.21 | 0.05 | 0 | 0.03 | 0.97 |
| naive | 69.27 | 0 | 0 | 1 | NA | 69.27 | 0 | 0 | 1 | NA |
| onlyT | 90 | 0 | 0 | 0.18 | 0.82 | 90 | 0 | 0 | 0.18 | 0.82 |

Table S13

**Decomposition of MSE into contributions from bias², SDSD, and LCS for Apache evaluation data BBCH30. Models are sorted in order of increasing RMSE for the Apache evaluation data averaged over stages (last column of Table 3). Due to rounding errors, the sum of fractions from bias², SDSD and LCS may not be exactly 1.00.**

|  | Apache evaluation data BBCH30 | | | | | | | | | |
| --- | --- | --- | --- | --- | --- | --- | --- | --- | --- | --- |
|  | Usual calibration | | | | | Protocol calibration | | | | |
| model | MSE | bias² | fraction  from bias² | fraction  from SDSD | fraction  from LCS | _MSE | bias² | fraction  from bias² | fraction  from SDSD | fraction  from LCS |
| M7 | 51.5 | 18.06 | 0.35 | 0.27 | 0.38 | 7.88 | 0.02 | 0 | 0.39 | 0.61 |
| M8 | 36.88 | 28.89 | 0.78 | 0.06 | 0.15 | 11.5 | 0.56 | 0.05 | 0.4 | 0.55 |
| e-median | 11.09 | 0.32 | 0.03 | 0.14 | 0.83 | 11.75 | 1.27 | 0.11 | 0.34 | 0.56 |
| e-mean | 12.3 | 2.32 | 0.19 | 0.26 | 0.56 | 15.17 | 2.23 | 0.15 | 0.22 | 0.63 |
| M20 | 62.88 | 31.64 | 0.5 | 0.22 | 0.27 | 14.88 | 1.27 | 0.09 | 0.39 | 0.53 |
| M12 | 17 | 5.06 | 0.3 | 0.09 | 0.62 | 17.12 | 3.52 | 0.21 | 0.26 | 0.53 |
| M16 | 25.25 | 0.25 | 0.01 | 0.37 | 0.62 | 16.75 | 0.25 | 0.01 | 0.46 | 0.52 |
| M13 | 21.25 | 5.06 | 0.24 | 0.24 | 0.52 | 13.5 | 0.06 | 0 | 0.64 | 0.35 |
| M17 | 25.88 | 3.52 | 0.14 | 0.07 | 0.8 | 35.75 | 14.06 | 0.39 | 0.07 | 0.53 |
| M15 | 34.12 | 4.52 | 0.13 | 0.12 | 0.74 | 29.62 | 4.52 | 0.15 | 0.1 | 0.75 |
| M25 | 26 | 12.25 | 0.47 | 0.14 | 0.39 | 13.25 | 4 | 0.3 | 0.3 | 0.39 |
| M26 | 42.38 | 11.39 | 0.27 | 0.31 | 0.43 | 44.12 | 11.39 | 0.26 | 0.24 | 0.5 |
| M21 | 33.75 | 18.06 | 0.54 | 0.07 | 0.39 | 36 | 18.06 | 0.5 | 0.08 | 0.41 |
| M24 | 19.5 | 0.25 | 0.01 | 0.04 | 0.95 | 32.88 | 17.02 | 0.52 | 0.03 | 0.45 |
| M9 | NA | NA | NA | NA | NA | NA | NA | NA | NA | NA |
| M2 | 20.62 | 4.52 | 0.22 | 0.11 | 0.67 | 120.88 | 87.89 | 0.73 | 0.02 | 0.26 |
| M30 | 148.62 | 19.14 | 0.13 | 0.18 | 0.69 | 143.88 | 87.89 | 0.61 | 0.03 | 0.36 |
| M1 | 99.62 | 40.64 | 0.41 | 0.13 | 0.46 | 106 | 36 | 0.34 | 0.17 | 0.49 |
| M6 | 168.62 | 92.64 | 0.55 | 0.29 | 0.16 | 102.62 | 47.27 | 0.46 | 0.31 | 0.23 |
| onlyT | 176.38 | 102.52 | 0.58 | 0.14 | 0.28 | 176.38 | 102.52 | 0.58 | 0.14 | 0.28 |
| naive | 187.4 | 96.46 | 0.51 | 0.49 | NA | 187.4 | 96.46 | 0.51 | 0.49 | NA |

Table S14

**Decomposition of MSE into contributions from bias², SDSD, and LCS for Apache evaluation data BBCH55. Models are sorted in order of increasing RMSE for the Apache evaluation data averaged over stages (last column of Table 3). Due to rounding errors, the sum of fractions from bias², SDSD and LCS may not be exactly 1.00.**

|  | Apache evaluation data BBCH55 | | | | | | | | | |
| --- | --- | --- | --- | --- | --- | --- | --- | --- | --- | --- |
|  | Usual calibration | | | | | Protocol calibration | | | | |
| model | MSE | bias² | fraction  from bias² | fraction  from SDSD | fraction  from LCS | _MSE | bias² | fraction  from bias² | fraction  from SDSD | fraction  from LCS |
| M7 | 14.75 | 4 | 0.27 | 0.02 | 0.71 | 27.25 | 0.06 | 0 | 0.12 | 0.88 |
| M8 | 14.5 | 3.06 | 0.21 | 0.06 | 0.73 | 22.75 | 0.06 | 0 | 0.15 | 0.85 |
| e-median | 19.12 | 4.52 | 0.24 | 0.07 | 0.69 | 23.38 | 2.64 | 0.11 | 0.11 | 0.77 |
| e-mean | 22.43 | 1.56 | 0.07 | 0.06 | 0.87 | 21.59 | 1.75 | 0.08 | 0.09 | 0.83 |
| M20 | 41.25 | 9 | 0.22 | 0.17 | 0.61 | 26 | 1 | 0.04 | 0.22 | 0.74 |
| M12 | 102 | 81 | 0.79 | 0.01 | 0.19 | 30.25 | 3.06 | 0.1 | 0.11 | 0.79 |
| M16 | 75.62 | 37.52 | 0.5 | 0.05 | 0.46 | 32 | 12.25 | 0.38 | 0.06 | 0.56 |
| M13 | 46.12 | 21.39 | 0.46 | 0.08 | 0.46 | 37.75 | 0.06 | 0 | 0.14 | 0.86 |
| M17 | 17 | 12.25 | 0.72 | 0.01 | 0.27 | 14.88 | 9.77 | 0.66 | 0.03 | 0.32 |
| M15 | 23 | 18.06 | 0.79 | 0.02 | 0.19 | 20.88 | 17.02 | 0.82 | 0.03 | 0.16 |
| M25 | 33.88 | 6.89 | 0.2 | 0.08 | 0.71 | 48.12 | 21.39 | 0.44 | 0.05 | 0.51 |
| M26 | 20.62 | 3.52 | 0.17 | 0.18 | 0.65 | 24.88 | 8.27 | 0.33 | 0.12 | 0.55 |
| M21 | 43.38 | 34.52 | 0.8 | 0 | 0.2 | 32.38 | 26.27 | 0.81 | 0.01 | 0.18 |
| M24 | 68.38 | 43.89 | 0.64 | 0.01 | 0.35 | 37.62 | 11.39 | 0.3 | 0.03 | 0.67 |
| M9 | 100.75 | 18.06 | 0.18 | 0.1 | 0.72 | 47.5 | 4 | 0.08 | 0.07 | 0.84 |
| M2 | 19.5 | 5.06 | 0.26 | 0.06 | 0.68 | 13.12 | 3.52 | 0.27 | 0.1 | 0.63 |
| M30 | 118.75 | 0.06 | 0 | 0.23 | 0.77 | 23.88 | 1.89 | 0.08 | 0.09 | 0.83 |
| M1 | 111.62 | 54.39 | 0.49 | 0.07 | 0.45 | 74.62 | 3.52 | 0.05 | 0.16 | 0.8 |
| M6 | 191.62 | 92.64 | 0.48 | 0.1 | 0.41 | 88.88 | 13.14 | 0.15 | 0.19 | 0.66 |
| onlyT | 123.5 | 52.56 | 0.43 | 0.07 | 0.5 | 123.5 | 52.56 | 0.43 | 0.07 | 0.5 |
| naive | 172.36 | 109.13 | 0.63 | 0.37 | NA | 172.36 | 109.13 | 0.63 | 0.37 | NA |

Table S15

**Decomposition of MSE into contributions from bias², SDSD, and LCS for Bermude evaluation data BBCH30. Models are sorted in order of increasing RMSE for the Bermude evaluation data averaged over stages (last column of Table 5). Due to rounding errors, the sum of fractions from bias², SDSD and LCS may not be exactly 1.00.**

|  | Bermude evaluation data BBCH30 | | | | | | | | | |
| --- | --- | --- | --- | --- | --- | --- | --- | --- | --- | --- |
|  | Usual calibration | | | | | Protocol calibration | | | | |
| model | MSE | bias² | fraction  from bias² | fraction  from SDSD | fraction  from LCS | _MSE | bias² | fraction  from bias² | fraction  from SDSD | fraction  from LCS |
| M15 | 25.25 | 0.56 | 0.02 | 0.36 | 0.61 | 30.5 | 2.25 | 0.07 | 0.37 | 0.56 |
| M2 | 49.38 | 17.02 | 0.34 | 0.22 | 0.44 | 20 | 4 | 0.2 | 0.28 | 0.52 |
| M17 | 23.38 | 0.77 | 0.03 | 0.29 | 0.68 | 48.5 | 20.25 | 0.42 | 0.2 | 0.38 |
| e-median | 42.44 | 5.06 | 0.12 | 0.25 | 0.63 | 24.19 | 0.02 | 0 | 0.38 | 0.62 |
| M8 | 164.5 | 105.06 | 0.64 | 0.13 | 0.23 | 26.12 | 3.52 | 0.13 | 0.44 | 0.43 |
| M21 | 42.5 | 14.06 | 0.33 | 0.17 | 0.49 | 47.62 | 17.02 | 0.36 | 0.19 | 0.45 |
| M12 | 57.5 | 33.06 | 0.57 | 0.08 | 0.34 | 21 | 0 | 0 | 0.26 | 0.74 |
| e-mean | 45.13 | 10.21 | 0.23 | 0.25 | 0.52 | 30.12 | 0.02 | 0 | 0.31 | 0.69 |
| M7 | 197.5 | 81 | 0.41 | 0.23 | 0.36 | 25.5 | 0.25 | 0.01 | 0.36 | 0.63 |
| M25 | 32.5 | 9 | 0.28 | 0.22 | 0.5 | 23.25 | 5.06 | 0.22 | 0.28 | 0.5 |
| M16 | 60.38 | 2.64 | 0.04 | 0.32 | 0.63 | 45.62 | 8.27 | 0.18 | 0.25 | 0.57 |
| M13 | 29.12 | 1.89 | 0.06 | 0.5 | 0.43 | 36.12 | 2.64 | 0.07 | 0.39 | 0.53 |
| M26 | 79.12 | 8.27 | 0.1 | 0.44 | 0.45 | 75.12 | 11.39 | 0.15 | 0.36 | 0.49 |
| M20 | 97.88 | 37.52 | 0.38 | 0.26 | 0.36 | 44.75 | 0 | 0 | 0.45 | 0.55 |
| M24 | 22 | 5.06 | 0.23 | 0 | 0.77 | 39 | 22.56 | 0.58 | 0 | 0.42 |
| M9 | NA | NA | NA | NA | NA | NA | NA | NA | NA | NA |
| M30 | 128.62 | 31.64 | 0.25 | 0.19 | 0.56 | 196.88 | 118.27 | 0.6 | 0.07 | 0.32 |
| M1 | 151.12 | 47.27 | 0.31 | 0.16 | 0.53 | 178.12 | 66.02 | 0.37 | 0.15 | 0.48 |
| naive | 152.18 | 84.25 | 0.55 | 0.45 | NA | 152.18 | 84.25 | 0.55 | 0.45 | NA |
| onlyT | 228.62 | 118.27 | 0.52 | 0.18 | 0.3 | 228.62 | 118.27 | 0.52 | 0.18 | 0.3 |
| M6 | 294.12 | 165.77 | 0.56 | 0.23 | 0.21 | 243.25 | 150.06 | 0.62 | 0.18 | 0.21 |

Table S16

**Decomposition of MSE into contributions from bias², SDSD, and LCS for Bermude evaluation data BBCH55. Models are sorted in order of increasing RMSE for the Bermude evaluation data averaged over stages (last column of Table 5). Due to rounding errors, the sum of fractions from bias², SDSD and LCS may not be exactly 1.00.**

|  | Bermude evaluation data BBCH55 | | | | | | | | | |
| --- | --- | --- | --- | --- | --- | --- | --- | --- | --- | --- |
|  | Usual calibration | | | | | Protocol calibration | | | | |
| model | MSE | bias² | fraction  from bias² | fraction  from SDSD | fraction  from LCS | _MSE | bias² | fraction  from bias² | fraction  from SDSD | fraction  from LCS |
| M15 | 8.5 | 0.25 | 0.03 | 0.02 | 0.95 | 11.25 | 0.56 | 0.05 | 0.04 | 0.91 |
| M2 | 19 | 0.25 | 0.01 | 0.03 | 0.96 | 21.25 | 4 | 0.19 | 0.01 | 0.8 |
| M17 | 12.88 | 6.89 | 0.54 | 0.01 | 0.46 | 6.38 | 0.77 | 0.12 | 0.07 | 0.81 |
| e-median | 15.5 | 0 | 0 | 0.02 | 0.98 | 22.12 | 2.64 | 0.12 | 0.03 | 0.85 |
| M8 | 10 | 0.06 | 0.01 | 0.01 | 0.98 | 22 | 3.06 | 0.14 | 0.07 | 0.79 |
| M21 | 20.88 | 11.39 | 0.55 | 0.01 | 0.45 | 9.75 | 4 | 0.41 | 0.02 | 0.57 |
| M12 | 121.88 | 97.52 | 0.8 | 0 | 0.2 | 32.5 | 3.06 | 0.09 | 0.01 | 0.89 |
| e-mean | 25.38 | 0 | 0 | 0.02 | 0.98 | 25.9 | 2.21 | 0.09 | 0.02 | 0.89 |
| M7 | 8.62 | 0.02 | 0 | 0 | 1 | 35 | 7.56 | 0.22 | 0.02 | 0.76 |
| M25 | 50.25 | 20.25 | 0.4 | 0.01 | 0.59 | 42 | 7.56 | 0.18 | 0.02 | 0.8 |
| M16 | 89.12 | 50.77 | 0.57 | 0.02 | 0.41 | 34 | 10.56 | 0.31 | 0.02 | 0.67 |
| M13 | 27 | 1.56 | 0.06 | 0.06 | 0.89 | 45.5 | 2.25 | 0.05 | 0.05 | 0.9 |
| M26 | 25.12 | 2.64 | 0.11 | 0.1 | 0.8 | 19.25 | 0.56 | 0.03 | 0.13 | 0.84 |
| M20 | 47.62 | 13.14 | 0.28 | 0.07 | 0.65 | 46.12 | 21.39 | 0.46 | 0.04 | 0.5 |
| M24 | 67.38 | 31.64 | 0.47 | 0.01 | 0.52 | 59.38 | 26.27 | 0.44 | 0 | 0.55 |
| M9 | 118.38 | 21.39 | 0.18 | 0.04 | 0.78 | 78.62 | 15.02 | 0.19 | 0.01 | 0.8 |
| M30 | 55.62 | 0.14 | 0 | 0.07 | 0.93 | 22.75 | 4 | 0.18 | 0.04 | 0.79 |
| M1 | 85.88 | 15.02 | 0.17 | 0.05 | 0.77 | 101.75 | 25 | 0.25 | 0.05 | 0.71 |
| naive | 132.65 | 59.79 | 0.45 | 0.55 | NA | 132.65 | 59.79 | 0.45 | 0.55 | NA |
| onlyT | 161.38 | 83.27 | 0.52 | 0.03 | 0.46 | 161.38 | 83.27 | 0.52 | 0.03 | 0.46 |
| M6 | 219.38 | 102.52 | 0.47 | 0.07 | 0.46 | 160 | 100 | 0.62 | 0.04 | 0.34 |

Table S17

**Decomposition of MSE into contributions from bias², SDSD, and LCS Australian calibration data BBCH30. Models are sorted in order of increasing RMSE for the Australian evaluation data averaged over stages (last column of Table 7). Due to rounding errors, the sum of fractions from bias², SDSD and LCS may not be exactly 1.00.**

|  | Australian calibration data BBCH30 | | | | | | | | | |
| --- | --- | --- | --- | --- | --- | --- | --- | --- | --- | --- |
|  | Usual calibration | | | | | Protocol calibration | | | | |
| model | MSE | bias² | fraction  from bias² | fraction  from SDSD | fraction  from LCS | _MSE | bias² | fraction  from bias² | fraction  from SDSD | fraction  from LCS |
| M9 | NA | NA | NA | NA | NA | NA | NA | NA | NA | NA |
| M17 | 71.64 | 6.75 | 0.09 | 0.02 | 0.89 | 63.24 | 0.02 | 0 | 0.02 | 0.98 |
| M13 | 201.05 | 47.4 | 0.24 | 0.13 | 0.63 | 113.59 | 1.72 | 0.02 | 0.11 | 0.88 |
| e-median | 85.62 | 1.16 | 0.01 | 0.22 | 0.76 | 97.93 | 0.13 | 0 | 0.16 | 0.84 |
| e-mean | 96.57 | 1.14 | 0.01 | 0.16 | 0.83 | 100.7 | 0.4 | 0 | 0.15 | 0.85 |
| M20 | NA | NA | NA | NA | NA | NA | NA | NA | NA | NA |
| M15 | 148.06 | 0.87 | 0.01 | 0.38 | 0.61 | 178.68 | 73.12 | 0.41 | 0.26 | 0.33 |
| M2 | 134.3 | 1.58 | 0.01 | 0.06 | 0.93 | 158.33 | 25.68 | 0.16 | 0.02 | 0.82 |
| M24 | 95.35 | 0 | 0 | 0.07 | 0.93 | 93.94 | 0.16 | 0 | 0.07 | 0.93 |
| M7 | 389.13 | 206.23 | 0.53 | 0.09 | 0.38 | 151.23 | 0.03 | 0 | 0.16 | 0.84 |
| M30 | 41.47 | 1.98 | 0.05 | 0 | 0.95 | 84.29 | 0.61 | 0.01 | 0.1 | 0.9 |
| M21 | 77.31 | 0.48 | 0.01 | 0.16 | 0.83 | 80.8 | 0 | 0 | 0.13 | 0.87 |
| M12 | 194.32 | 21.97 | 0.11 | 0.03 | 0.86 | 164.62 | 1.04 | 0.01 | 0.01 | 0.98 |
| M28 | 599.03 | 11.25 | 0.02 | 0.47 | 0.51 | 151.54 | 0.55 | 0 | 0 | 1 |
| M25 | 143.35 | 12.22 | 0.09 | 0.01 | 0.9 | 152.5 | 21.97 | 0.14 | 0.03 | 0.82 |
| M8 | 273.19 | 106.36 | 0.39 | 0.13 | 0.48 | 131.63 | 0.47 | 0 | 0.01 | 0.98 |
| M26 | 212.1 | 52.79 | 0.25 | 0.02 | 0.73 | 347.35 | 135.64 | 0.39 | 0 | 0.61 |
| M16 | 193.63 | 32.88 | 0.17 | 0.55 | 0.28 | 150.94 | 0 | 0 | 0.46 | 0.54 |
| M1 | 291.27 | 115.23 | 0.4 | 0 | 0.6 | 178.32 | 0.25 | 0 | 0.01 | 0.99 |
| naive | 220.81 | 0 | 0 | 1 | NA | 220.81 | 0 | 0 | 1 | NA |
| M29 | 163.25 | 0.21 | 0 | 0.15 | 0.85 | 201.77 | 4.31 | 0.02 | 0 | 0.98 |
| onlyT | 168.59 | 0.01 | 0 | 0 | 1 | 168.59 | 0.01 | 0 | 0 | 1 |
| M6 | 108.62 | 6.27 | 0.06 | 0.04 | 0.9 | 129.81 | 0.09 | 0 | 0.06 | 0.94 |

Table S18

**Decomposition of MSE into contributions from bias², SDSD, and LCS Australian calibration data BBCH65. Models are sorted in order of increasing RMSE for the Australian evaluation data averaged over stages (last column of Table 7). Due to rounding errors, the sum of fractions from bias², SDSD and LCS may not be exactly 1.00.**

|  | Australian calibration data BBCH65 | | | | | | | | | |
| --- | --- | --- | --- | --- | --- | --- | --- | --- | --- | --- |
|  | Usual calibration | | | | | Protocol calibration | | | | |
| model | MSE | bias² | fraction  from bias² | fraction  from SDSD | fraction  from LCS | _MSE | bias² | fraction  from bias² | fraction  from SDSD | fraction  from LCS |
| M9 | 31.81 | 0.25 | 0.01 | 0.06 | 0.93 | 38.63 | 0.08 | 0 | 0.49 | 0.51 |
| M17 | 73.97 | 0.16 | 0 | 0.32 | 0.68 | 82.57 | 13.37 | 0.16 | 0.27 | 0.56 |
| M13 | 103.38 | 1.94 | 0.02 | 0.46 | 0.52 | 75.46 | 3.11 | 0.04 | 0.4 | 0.56 |
| e-median | 32.72 | 1.94 | 0.06 | 0.5 | 0.44 | 38.95 | 0.85 | 0.02 | 0.49 | 0.49 |
| e-mean | 33.68 | 0.88 | 0.03 | 0.44 | 0.53 | 37.32 | 0.83 | 0.02 | 0.51 | 0.47 |
| M20 | 94.54 | 20.74 | 0.22 | 0.47 | 0.31 | 73.88 | 8.53 | 0.12 | 0.23 | 0.65 |
| M15 | 176.38 | 0.85 | 0 | 0.43 | 0.56 | 101.2 | 7.93 | 0.08 | 0.35 | 0.57 |
| M2 | 33.8 | 4.54 | 0.13 | 0.35 | 0.52 | 32.51 | 5 | 0.15 | 0.32 | 0.53 |
| M24 | 95.96 | 4.33 | 0.05 | 0.53 | 0.43 | 89.77 | 0.06 | 0 | 0.58 | 0.42 |
| M7 | 574.03 | 282.74 | 0.49 | 0.19 | 0.32 | 100.35 | 0.06 | 0 | 0.45 | 0.55 |
| M30 | 144.34 | 53.89 | 0.37 | 0.1 | 0.53 | 124.1 | 90.27 | 0.73 | 0.09 | 0.18 |
| M21 | 53.51 | 0 | 0 | 0.55 | 0.45 | 30.1 | 0.06 | 0 | 0.35 | 0.65 |
| M12 | 72.13 | 2.75 | 0.04 | 0.07 | 0.9 | 46.67 | 0.02 | 0 | 0.01 | 0.99 |
| M28 | 511.02 | 143.39 | 0.28 | 0.66 | 0.06 | 40.31 | 0.02 | 0 | 0.11 | 0.89 |
| M25 | 25.61 | 0.2 | 0.01 | 0.02 | 0.97 | 40.89 | 15.38 | 0.38 | 0.1 | 0.53 |
| M8 | 444.19 | 243.49 | 0.55 | 0.13 | 0.32 | 65.23 | 0.08 | 0 | 0.29 | 0.71 |
| M26 | 74.25 | 42.26 | 0.57 | 0.03 | 0.41 | 40.42 | 0 | 0 | 0.1 | 0.9 |
| M16 | 215.19 | 0.01 | 0 | 0.73 | 0.27 | 149.29 | 0.58 | 0 | 0.67 | 0.32 |
| M1 | 42.21 | 1.4 | 0.03 | 0.19 | 0.78 | 56.54 | 10.48 | 0.19 | 0.15 | 0.66 |
| naive | 590.17 | 0 | 0 | 1 | NA | 590.17 | 0 | 0 | 1 | NA |
| M29 | 169.43 | 43.64 | 0.26 | 0.09 | 0.65 | 80.86 | 4.77 | 0.06 | 0.04 | 0.9 |
| onlyT | 236.45 | 0.58 | 0 | 0.05 | 0.94 | 236.45 | 0.58 | 0 | 0.05 | 0.94 |

Table S19

**Decomposition of MSE into contributions from bias², SDSD, and LCS Australian calibration data BBCH90. Models are sorted in order of increasing RMSE for the Australian evaluation data averaged over stages (last column of Table 7). Due to rounding errors, the sum of fractions from bias², SDSD and LCS may not be exactly 1.00.**

|  | Australian calibration data BBCH90 | | | | | | | | | |
| --- | --- | --- | --- | --- | --- | --- | --- | --- | --- | --- |
|  | Usual calibration | | | | | Protocol calibration | | | | |
| model | MSE | bias² | fraction  from bias² | fraction  from SDSD | fraction  from LCS | _MSE | bias² | fraction  from bias² | fraction  from SDSD | fraction  from LCS |
| M9 | 125.64 | 0.04 | 0 | 0 | 1 | 57.68 | 0.36 | 0.01 | 0.08 | 0.92 |
| M17 | 315.62 | 297.56 | 0.94 | 0.06 | 0 | 8.12 | 5.06 | 0.62 | 0.38 | 0 |
| M13 | 170.32 | 4.82 | 0.03 | 0.03 | 0.94 | 53.39 | 0.65 | 0.01 | 0.03 | 0.96 |
| e-median | 52.18 | 1.31 | 0.03 | 0.01 | 0.96 | 53.59 | 0.01 | 0 | 0.07 | 0.93 |
| e-mean | 70.94 | 0.58 | 0.01 | 0 | 0.99 | 60.38 | 0.8 | 0.01 | 0.04 | 0.95 |
| M20 | 146.81 | 14.41 | 0.1 | 0.19 | 0.71 | 123.67 | 0.63 | 0.01 | 0 | 0.99 |
| M15 | 8.98 | 0.01 | 0 | 0.24 | 0.76 | 9.56 | 0.36 | 0.04 | 0.05 | 0.91 |
| M2 | 88.16 | 2.55 | 0.03 | 0.02 | 0.95 | 84.3 | 0.48 | 0.01 | 0.02 | 0.98 |
| M24 | 159.83 | 0.5 | 0 | 0.02 | 0.98 | 202.73 | 53.35 | 0.26 | 0.01 | 0.73 |
| M7 | 172.68 | 11.53 | 0.07 | 0.02 | 0.92 | 64.49 | 0.16 | 0 | 0.1 | 0.89 |
| M30 | 0.1 | 0 | 0 | 0.13 | 0.87 | 77.53 | 26.05 | 0.34 | 0.1 | 0.57 |
| M21 | 40.69 | 5.27 | 0.13 | 0.25 | 0.62 | 60.6 | 0.04 | 0 | 0.08 | 0.92 |
| M12 | 80.79 | 37.16 | 0.46 | 0.02 | 0.52 | 41.87 | 16.84 | 0.4 | 0 | 0.6 |
| M28 | 422.48 | 163.94 | 0.39 | 0.28 | 0.33 | 47.51 | 1.01 | 0.02 | 0.01 | 0.97 |
| M25 | 57.1 | 0.16 | 0 | 0.03 | 0.97 | 71.67 | 13.66 | 0.19 | 0 | 0.81 |
| M8 | 193.05 | 67.17 | 0.35 | 0 | 0.65 | 55.2 | 0.25 | 0 | 0.06 | 0.93 |
| M26 | 39.69 | 0.09 | 0 | 0.01 | 0.98 | 89.03 | 28.05 | 0.32 | 0.25 | 0.44 |
| M16 | 544.76 | 397.71 | 0.73 | 0.12 | 0.15 | 310.3 | 96.73 | 0.31 | 0.22 | 0.46 |
| M1 | 532.83 | 338.41 | 0.64 | 0 | 0.36 | 360.19 | 171.51 | 0.48 | 0.01 | 0.51 |
| naive | 518.97 | 0 | 0 | 1 | NA | 518.97 | 0 | 0 | 1 | NA |
| M29 | NA | NA | NA | NA | NA | NA | NA | NA | NA | NA |

Table S20

**Decomposition of MSE into contributions from bias², SDSD, and LCS Australian evaluation data BBCH30. Models are sorted in order of increasing RMSE for the Australian evaluation data averaged over stages (last column of Table 7). Due to rounding errors, the sum of fractions from bias², SDSD and LCS may not be exactly 1.00.**

|  | Australian evaluation data BBCH30 | | | | | | | | | |
| --- | --- | --- | --- | --- | --- | --- | --- | --- | --- | --- |
|  | Usual calibration | | | | | Protocol calibration | | | | |
| model | MSE | bias² | fraction  from bias² | fraction  from SDSD | fraction  from LCS | _MSE | bias² | fraction  from bias² | fraction  from SDSD | fraction  from LCS |
| M9 | NA | NA | NA | NA | NA | NA | NA | NA | NA | NA |
| M17 | 118.41 | 6.75 | 0.06 | 0.02 | 0.92 | 146.49 | 27.94 | 0.19 | 0.03 | 0.78 |
| M13 | 198.66 | 31.94 | 0.16 | 0.48 | 0.36 | 135.96 | 3.89 | 0.03 | 0.19 | 0.78 |
| e-median | 142.51 | 0.1 | 0 | 0.16 | 0.84 | 148.47 | 16.03 | 0.11 | 0.13 | 0.76 |
| e-mean | 130.24 | 3 | 0.02 | 0.18 | 0.8 | 157.77 | 16.73 | 0.11 | 0.12 | 0.77 |
| M20 | NA | NA | NA | NA | NA | NA | NA | NA | NA | NA |
| M15 | 157.44 | 4.9 | 0.03 | 0.09 | 0.88 | 183.62 | 46.78 | 0.25 | 0.15 | 0.59 |
| M2 | 201.47 | 25.36 | 0.13 | 0.03 | 0.84 | 270.99 | 94.54 | 0.35 | 0.02 | 0.63 |
| M24 | 146.58 | 2.55 | 0.02 | 0.06 | 0.92 | 148.03 | 4.4 | 0.03 | 0.06 | 0.91 |
| M7 | 370.27 | 200.28 | 0.54 | 0.24 | 0.22 | 202.82 | 10.8 | 0.05 | 0.12 | 0.82 |
| M30 | 116.36 | 7.71 | 0.07 | 0.1 | 0.83 | 114.08 | 3.65 | 0.03 | 0.07 | 0.9 |
| M21 | 146.29 | 22.9 | 0.16 | 0 | 0.84 | 231.21 | 71.79 | 0.31 | 0.01 | 0.68 |
| M12 | 241.14 | 44.36 | 0.18 | 0.02 | 0.8 | 197.11 | 7.08 | 0.04 | 0 | 0.96 |
| M28 | 651.99 | 303.13 | 0.46 | 0.17 | 0.36 | 226.73 | 32.04 | 0.14 | 0 | 0.85 |
| M25 | 173.13 | 32.75 | 0.19 | 0.03 | 0.78 | 234.63 | 71.79 | 0.31 | 0.01 | 0.68 |
| M8 | 256.1 | 58.55 | 0.23 | 0.06 | 0.72 | 377.55 | 128.78 | 0.34 | 0 | 0.66 |
| M26 | 169.71 | 21.64 | 0.13 | 0.09 | 0.78 | 279.84 | 47.64 | 0.17 | 0 | 0.83 |
| M16 | 263.87 | 38.73 | 0.15 | 0.44 | 0.41 | 244.36 | 5.81 | 0.02 | 0.65 | 0.33 |
| M1 | 479.88 | 277.58 | 0.58 | 0.06 | 0.36 | 241 | 35.68 | 0.15 | 0.14 | 0.71 |
| naive | 280.57 | 4.38 | 0.02 | 0.98 | NA | 280.57 | 4.38 | 0.02 | 0.98 | NA |
| M29 | 160.01 | 3.89 | 0.02 | 0.26 | 0.72 | 372.11 | 124.56 | 0.33 | 0.04 | 0.63 |

Table S21

**Decomposition of MSE into contributions from bias², SDSD, and LCS Australian evaluation data BBCH65. Models are sorted in order of increasing RMSE for the Australian evaluation data averaged over stages (last column of Table 7). Due to rounding errors, the sum of fractions from bias², SDSD and LCS may not be exactly 1.00.**

|  | Australian evaluation data BBCH65 | | | | | | | | | |
| --- | --- | --- | --- | --- | --- | --- | --- | --- | --- | --- |
|  | Usual calibration | | | | | Protocol calibration | | | | |
| model | MSE | bias² | fraction  from bias² | fraction  from SDSD | fraction  from LCS | _MSE | bias² | fraction  from bias² | fraction  from SDSD | fraction  from LCS |
| M9 | 121.64 | 43.18 | 0.35 | 0.01 | 0.63 | 103.46 | 41.02 | 0.4 | 0.11 | 0.5 |
| M17 | 88.83 | 46.15 | 0.52 | 0 | 0.48 | 51.12 | 10.13 | 0.2 | 0 | 0.8 |
| M13 | 91.38 | 22.98 | 0.25 | 0.19 | 0.56 | 94.66 | 27.44 | 0.29 | 0.04 | 0.67 |
| e-median | 71.9 | 21.41 | 0.3 | 0.03 | 0.67 | 93.54 | 36.52 | 0.39 | 0.02 | 0.59 |
| e-mean | 70.97 | 22.53 | 0.32 | 0.03 | 0.66 | 92.06 | 44.92 | 0.49 | 0.02 | 0.49 |
| M20 | 192.22 | 95.91 | 0.5 | 0.04 | 0.46 | 51.14 | 3.72 | 0.07 | 0.03 | 0.9 |
| M15 | 44.02 | 4.52 | 0.1 | 0 | 0.9 | 74.84 | 41.02 | 0.55 | 0 | 0.45 |
| M2 | 75.73 | 11.59 | 0.15 | 0 | 0.84 | 72.15 | 12.36 | 0.17 | 0 | 0.83 |
| M24 | 88.79 | 40.31 | 0.45 | 0.02 | 0.52 | 67.35 | 19.4 | 0.29 | 0.03 | 0.68 |
| M7 | 133.98 | 57.69 | 0.43 | 0.15 | 0.41 | 122.27 | 38.91 | 0.32 | 0.03 | 0.65 |
| M30 | 64.32 | 0.35 | 0.01 | 0.01 | 0.99 | 221.94 | 159.44 | 0.72 | 0 | 0.28 |
| M21 | 71.08 | 36.19 | 0.51 | 0 | 0.49 | 149.59 | 91.61 | 0.61 | 0.01 | 0.38 |
| M12 | 87.54 | 0.12 | 0 | 0 | 1 | 77.82 | 0.39 | 0.01 | 0.02 | 0.97 |
| M28 | 716.64 | 584.79 | 0.82 | 0.07 | 0.11 | 173.4 | 81.28 | 0.47 | 0.01 | 0.53 |
| M25 | 71.61 | 12.75 | 0.18 | 0.01 | 0.81 | 178.94 | 117.7 | 0.66 | 0 | 0.34 |
| M8 | 126.25 | 52.74 | 0.42 | 0.01 | 0.57 | 104.22 | 24.05 | 0.23 | 0.01 | 0.76 |
| M26 | 164.23 | 105.96 | 0.65 | 0.01 | 0.35 | 122.19 | 34.86 | 0.29 | 0.11 | 0.6 |
| M16 | 141.28 | 38.22 | 0.27 | 0.29 | 0.44 | 180.87 | 76.35 | 0.42 | 0.28 | 0.29 |
| M1 | 136.88 | 43.91 | 0.32 | 0.1 | 0.58 | 213.82 | 127.54 | 0.6 | 0.05 | 0.35 |
| naive | 299.01 | 24.28 | 0.08 | 0.92 | NA | 299.01 | 24.28 | 0.08 | 0.92 | NA |
| M29 | 93.77 | 49.22 | 0.52 | 0.08 | 0.39 | 100.32 | 4.29 | 0.04 | 0.08 | 0.88 |

Table S22

**Decomposition of MSE into contributions from bias², SDSD, and LCS Australian evaluation data BBCH90. Models are sorted in order of increasing RMSE for the Australian evaluation data averaged over stages (last column of Table 7). Due to rounding errors, the sum of fractions from bias², SDSD and LCS may not be exactly 1.00.**

|  | Australian evaluation data BBCH90 | | | | | | | | | |
| --- | --- | --- | --- | --- | --- | --- | --- | --- | --- | --- |
|  | Usual calibration | | | | | Protocol calibration | | | | |
| model | MSE | bias² | fraction  from bias² | fraction  from SDSD | fraction  from LCS | _MSE | bias² | fraction  from bias² | fraction  from SDSD | fraction  from LCS |
| M9 | 12 | 0.16 | 0.01 | 0.07 | 0.92 | 30 | 5.76 | 0.19 | 0.02 | 0.79 |
| M17 | 130 | 121 | 0.93 | 0.07 | 0 | 30.5 | 30.25 | 0.99 | 0.01 | 0 |
| M13 | 11.6 | 0.64 | 0.06 | 0 | 0.94 | 11.6 | 0.16 | 0.01 | 0.06 | 0.92 |
| e-median | 13.3 | 4 | 0.3 | 0.01 | 0.69 | 10.65 | 0.25 | 0.02 | 0 | 0.98 |
| e-mean | 9.99 | 1.13 | 0.11 | 0.03 | 0.86 | 11.16 | 0.95 | 0.08 | 0 | 0.92 |
| M20 | 80 | 51.84 | 0.65 | 0.01 | 0.35 | 120.4 | 12.96 | 0.11 | 0.01 | 0.88 |
| M15 | 40.4 | 10.24 | 0.25 | 0.21 | 0.53 | 35.6 | 7.84 | 0.22 | 0.2 | 0.58 |
| M2 | 12 | 2.56 | 0.21 | 0.03 | 0.76 | 11.4 | 1 | 0.09 | 0.08 | 0.83 |
| M24 | 19.2 | 2.56 | 0.13 | 0.29 | 0.57 | 69.4 | 54.76 | 0.79 | 0.06 | 0.15 |
| M7 | 14.6 | 1.96 | 0.13 | 0.01 | 0.86 | 15.2 | 0 | 0 | 0.04 | 0.96 |
| M30 | 20.2 | 11.56 | 0.57 | 0.01 | 0.42 | 13.8 | 1.96 | 0.14 | 0.04 | 0.82 |
| M21 | 26 | 12.96 | 0.5 | 0.06 | 0.44 | 7.8 | 0.04 | 0.01 | 0.07 | 0.93 |
| M12 | 86.4 | 57.76 | 0.67 | 0.05 | 0.28 | 75.2 | 16 | 0.21 | 0.11 | 0.67 |
| M28 | 334.6 | 302.76 | 0.9 | 0.03 | 0.06 | 13.4 | 1.96 | 0.15 | 0.23 | 0.62 |
| M25 | 27.2 | 7.84 | 0.29 | 0.17 | 0.54 | 18.8 | 10.24 | 0.54 | 0.01 | 0.45 |
| M8 | 55.6 | 36 | 0.65 | 0.03 | 0.32 | 16.4 | 0.16 | 0.01 | 0.03 | 0.96 |
| M26 | 26.2 | 0.36 | 0.01 | 0.37 | 0.62 | 46.6 | 17.64 | 0.38 | 0.2 | 0.42 |
| M16 | 468.8 | 432.64 | 0.92 | 0 | 0.08 | 60.6 | 38.44 | 0.63 | 0.19 | 0.17 |
| M1 | 285.6 | 268.96 | 0.94 | 0.01 | 0.05 | 143.2 | 125.44 | 0.88 | 0.07 | 0.05 |
| naive | 97.85 | 0.01 | 0 | 1 | NA | 97.85 | 0.01 | 0 | 1 | NA |
| M29 | NA | NA | NA | NA | NA | NA | NA | NA | NA | NA |

Table S23

The decomposition of MSE on the average over modeling teams and over BBCH30 and BBCH55 for the French data sets and over BBCH30, BBCH65, and BBCH90 for the Australian data set.

| Data set | calibration | Bias²/MSE | SDSD/MSE | LCS/MSE |
| --- | --- | --- | --- | --- |
| Apache calibration data | usual | 0.15 | 0.18 | 0.67 |
|  | protocol | 0.12 | 0.15 | 0.73 |
| Apache evaluation data | usual | 0.39 | 0.13 | 0.47 |
|  | protocol | 0.35 | 0.13 | 0.52 |
| Bermude calibration data | usual | 0.23 | 0.19 | 0.58 |
|  | protocol | 0.16 | 0.11 | 0.73 |
| Bermude evaluation data | usual | 0.38 | 0.15 | 0.47 |
|  | protocol | 0.36 | 0.14 | 0.50 |
| Australian calibration data | usual | 0.23 | 0.27 | 0.52 |
|  | protocol | 0.10 | 0.24 | 0.69 |
| Australian evaluation data | usual | 0.35 | 0.16 | 0.51 |
|  | protocol | 0.28 | 0.16 | 0.59 |

Table S24

**Comparison of the AICc and BIC criteria for model selection for the French datasets. Results are only shown for those modeling teams where AICc and BIC led to different final choices of estimated parameters. For each of those modeling teams, the final number of parameters (n) and RMSE for each stage using the evaluation data are shown.**

|  |  | AICc | | | BIC | | |
| --- | --- | --- | --- | --- | --- | --- | --- |
|  |  | n | RMSE BBCH30 | RMSE BBCH55 | n | RMSE BBCH30 | RMSE BBCH55 |
| Apache | M13 | 4 | 3.3 | 5.6 | 3 | 3.7 | 6.1 |
|  | M16 | 4 | 6.6 | 5.0 | 3 | 6.6 | 5.0 |
|  | average |  | 5.0 | 5.3 |  | 5.2 | 5.6 |
| Bermude | M20 | 5 | 6.9 | 6.3 | 4 | 6.7 | 6.8 |

Table S25

**Comparison of the AICc and BIC criteria for model selection for the Australian datasets. Results are only shown for those modeling teams where AICc and BIC led to different final models. For each of those modeling teams, the final number of parameters (n) and RMSE for each stage using the evaluation data are shown.**

|  | AICc | | | | BIC | | | |
| --- | --- | --- | --- | --- | --- | --- | --- | --- |
|  | n | RMSE BBCH30 | RMSE BBCH65 | RMSE BBCH90 | n | RMSE BBCH30 | RMSE BBCH65 | RMSEBBCH90 |
| M2 | 8 | 16.46 | 8.61 | 3.74 | 7 | 16.46 | 8.49 | 3.38 |
| M6 | 7 | 14.8 | 24.14 | 11.44 | 6 | 14.55 | 24.42 | 10.79 |
| M12 | 8 | 13.46 | 9.26 | 15.72 | 7 | 14.04 | 8.82 | 8.67 |
| M17 | 8 | 12.94 | 7.43 | 5.1 | 7 | 12.1 | 7.15 | 5.52 |
| M21 | 8 | 16.12 | 11.95 | 2.49 | 7 | 15.21 | 12.23 | 2.79 |
| average |  | 14.8 | 12.3 | 7.7 |  | 14.5 | 12.2 | 6.2 |
